# CryoENsemble - a Bayesian approach for reweighting biomolecular structural ensembles using heterogeneous cryo-EM maps

**DOI:** 10.1101/2023.11.21.567999

**Authors:** Tomasz Włodarski, Julian O. Streit, Alkistis Mitropoulou, Lisa D. Cabrita, Michele Vendruscolo, John Christodoulou

**Affiliations:** Institute of Structural and Molecular Biology, University College London, UK; Institute of Biochemistry and Biophysics, Polish Academy of Sciences, Warsaw, Poland; Birkbeck College, University of London, UK; Centre for Misfolding Diseases, Yusuf Hamied Department of Chemistry, University of Cambridge, Cambridge, UK

**Keywords:** Statistical inference, molecular dynamics simulations, cryo-EM, trigger factor, structural biology

## Abstract

Cryogenic electron microscopy (cryo-EM) has emerged as a central tool for the determination of structures of complex biological molecules. Accurately characterising the dynamics of such systems, however, remains a challenge. To address this, we introduce cryoENsemble, a method that applies Bayesian reweighing to conformational ensembles derived from molecular dynamics simulations to improve their agreement with cryo-EM data and extract dynamics information. We illustrate the use of cryoENsemble to determine the dynamics of the ribosome-bound state of the co-translational chaperone trigger factor (TF). We also show that cryoENsemble can assist with the interpretation of low-resolution, noisy or unaccounted regions of cryo-EM maps. Notably, we are able to link an unaccounted part of the cryo-EM map to the presence of another protein (methionine aminopeptidase, or MetAP), rather than to the dynamics of TF, and model its TF-bound state. Based on these results, cryoENsemble is expected to find use for challenging heterogeneous cryo-EM maps for various biomolecular systems, especially those encompassing dynamic elements.

## Introduction

Describing the dynamics of complex macromolecular systems presents significant challenges^1^. The main techniques to achieve this goal are nuclear magnetic resonance (NMR) spectroscopy and single-molecule fluorescence methods^2–5^. More recently, technological and methodological advancements in single-particle cryogenic electron microscopy (cryo-EM), including improvements in electron detectors, image processing software and motion correction algorithms, have offered a new means to investigate protein dynamics^6–9^. By the recording of millions two-dimensional (2D) projection images of biomolecules captured by flash freezing in various compositional or conformational states, cryo-EM offers a glimpse into the diverse conformational landscape of dynamic macromolecular complexes.

A variety of computational methods for fitting and refining atomic models against single-particle cryo-EM data have been developed^10–13^. These methods include rigid body fitting of available X-ray structures into low-resolution cryo-EM maps^14^, incorporation of protein flexibility through normal mode analysis^15^ and flexible fitting^16,17^, and density-based molecular dynamics (MD) simulations^18–21^. Despite these advances, however, characterising the conformational heterogeneity underlying the dynamics of the systems under observation in cryo-EM samples remains a significant challenge^22–24^. Structural regions that display conformational heterogeneity can be incorrectly aligned and then erroneously averaged with other images, causing these regions to become blurred, or even invisible, in the reconstruction and leading to lower final resolution and less detailed or incomplete maps. Separating these regions into homogeneous subsets during post-processing can be achieved, for example, by using heterogeneous refinement with maximum likelihood classification methods^25^. This approach, however, tends to work better for discrete heterogeneity, when the system can be characterised by a finite number of states. For continuous conformational heterogeneity, other methods have been developed, including focus refinement, where a mask is applied to different regions of the structure^26^, multi-body refinement^27^, manifold embedding^28^ or deep neural networks^29^.

Typically, to generate dynamical descriptions, structural models can be fine-tuned with experimental data^30–35^. This approach, however, presents significant challenges as the experimental data are affected by a combination of the experimental errors and approximations included in post-processing into molecular simulations. As a result, different conformations can lead to a similar agreement with experimental observables, particularly when the data are incomplete and noisy, or when the forward model is dependent on many approximations, such as being based only on distances or angles between atoms to back-calculate experimental properties. Solutions that combine structural information from various experimental techniques (e.g. NMR spectroscopy, cryo-EM, small-angle X-ray scattering (SAXS)^30^) with computational methods (e.g. molecular dynamics) and Bayesian inference have been proposed to produce structural ensembles^30–32^. This integrative structural biology approach^33^ has been applied to many biological systems^34,35^. Bayesian inference can be applied during MD simulations by adding a bias energy term to constrain simulations to sample conformations in agreement with experimental data^35^, or it can be applied *a posteriori* when the experimental data are used to reweight the MD ensemble. The utility of these methods depends on the nature of the system under study, as well as the available experimental data^36^.

Here, we describe cryoENsemble, a computational approach that combines molecular dynamics simulations with Bayesian reweighting utilising cryo-EM maps (**Fig. 1**). This method allows the interpretation of discrete and continuous heterogeneity from cryo-EM maps to describe the underlying structural ensembles accurately. To accomplish this, we adapted and extended the Bayesian Inference Of ENsembles (BioEn) method^37^ that uses various experimental data (e.g. NMR, SAXS, DEER) to refine structural ensembles from MD simulations. We first validated the cryoENsemble method with synthetic cryo-EM maps from two well-characterized systems, namely adenylate kinase (ADK) and ribosomal nascent chain complex (RNC) (**Fig. 2**). We could effectively reweight the structural ensembles in both cases, capture important structural features and, at the same time, account for variations in resolution and noise levels present in the density maps that correspond to discrete and continuous cryo-EM heterogeneity (see Methods).

**Figure 1.**
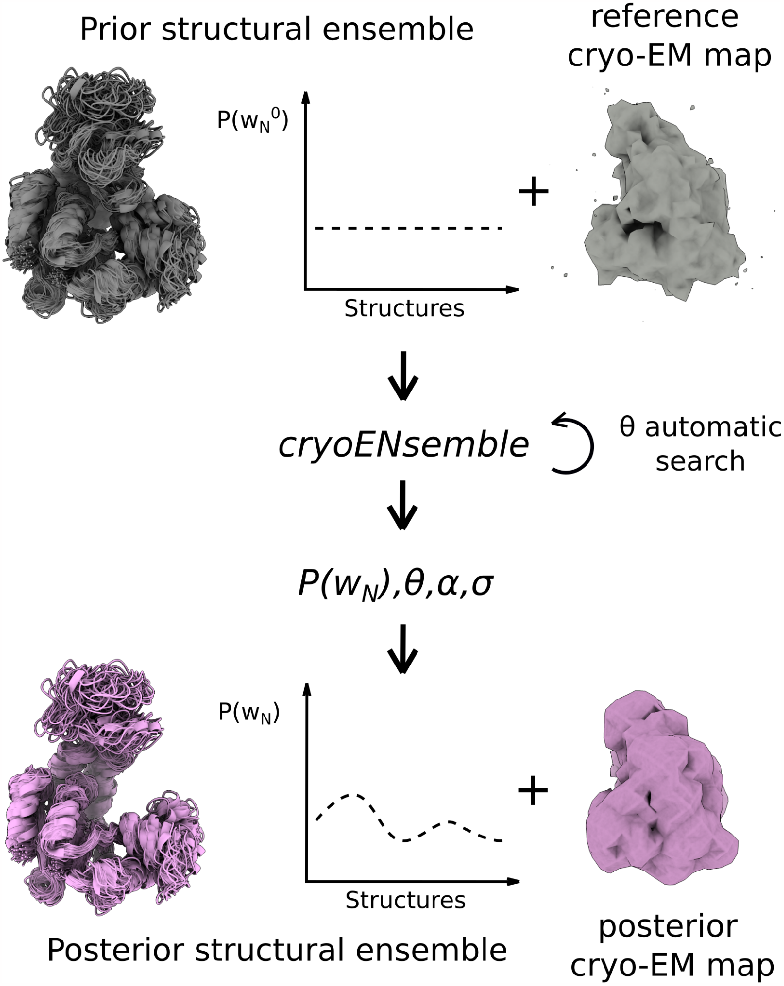
Schematic illustration of the cryoENsemble method. The input includes a structural ensemble (in grey), typically obtained from molecular dynamics simulations, and a cryo-EM map of the biological system under investigation. Each model from the structural ensemble is fitted into the density prior to the cryoENsemble calculations. The prior 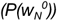 and posterior (P(w_*N*_)) structural ensembles consist of N structural models with 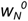 and w_*N*_ weights, respectively. The parameters θ, α and σ are obtained during the reweighting along with the posterior average cryo-EM map (in pink).

**Figure 2.**
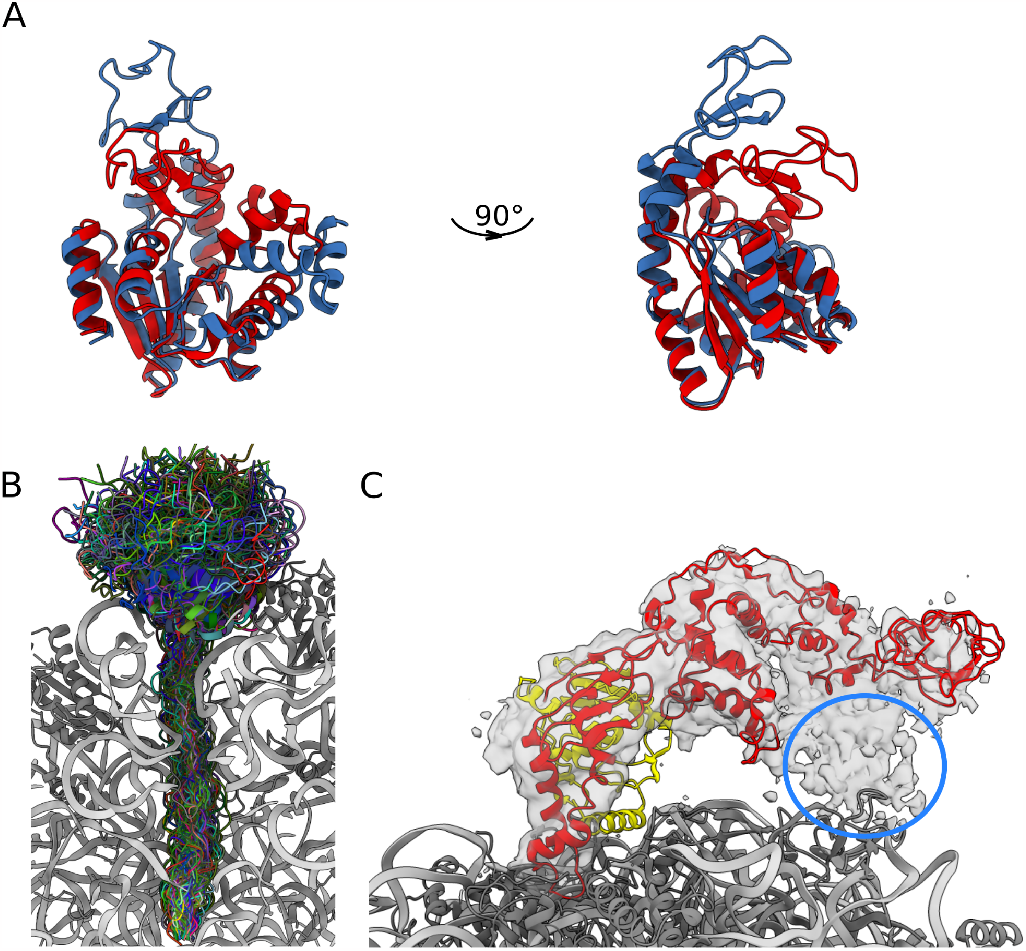
Molecular systems used for the development and testing of the cryoENsemble method. **(A)** X-ray structures of the E. coli adenylate kinase (ADK) in the open (PDB ID: 4AKE^48^, shown in blue) and closed (PDB ID: 1AKE^49^, shown in red) states. **(B)** Structural ensemble of FLN5-6 ribosome nascent chain complex (FLN5-6 RNC) consisting of 100 structures, randomly selected from the all-atom structure-based MD simulation^50^ that are used in the reweighting process. Each structure is depicted in a different colour and combines the N-terminal folded FLN5 domain followed by 31 amino acid linker consisting of the subsequent FLN6 domain and a SecM stalling sequence covalently bound to the tRNA at the peptidyl transferase centre. A cross-section of the 70S ribosome is shown for clarity (in grey ribosomal proteins and in silver rRNA). **(C)** Trigger factor (in red) and peptide deformylase (in yellow) bound to the 70S ribosome (ribosomal proteins in grey, rRNA in silver) (from PDB ID: 7D80^38^) with cryo-EM density corresponding to TF bound states. The region of incompletely characterised density is depicted by a blue circle (from EMDB: 30611^38^).

Next, we applied cryoENsemble to the cryo-EM map of the ribosome-bound state of trigger factor (TF) stabilised by the presence of peptide deformylase (PDF) and methionine aminopeptidase (MetAP) (**Fig. 2**). TF is the only ribosome-associated chaperone in bacteria, whereas PDF and MetAP are essential enzymes involved in the co-translational removal of formylated methionine in nascent protein chains, and both bind in the proximity of the ribosomal exit tunnel^38^. Despite intensive research^39–44^, the detailed role of TF in the co-translational folding process remains incompletely understood due to the experimental challenges presented by its dynamic nature, even in its ribosome-bound state, and only low-resolution or incomplete cryo-EM maps, that often encompass merely the ribosome-binding domain (RBD)^45–47^ are available.

By using all-atom MD simulations combined with cryoENsemble, we provide insights into the dynamics of TF, as captured within this cryo-EM map and explain the additional density present around TF. Our findings indicate that an ensemble of TF structures obtained with MD can better explain cryo-EM maps compared to a single model. Furthermore, using cryoENsemble, we confirmed that the additional density localised close to TF is not due to the dynamics of TF, as was initially hypothesised^45^. Instead, by fitting MetAP to this unaccounted density, we found a compelling overlap, further confirming that this density stems from a novel binding site of the MetAP, as suggested recently^38^.

Overall, we demonstrate that cryoENsemble can extract otherwise elusive information about macromolecular dynamics from heterogenous cryo-EM data. It also proves valuable in modelling biomolecular complexes, when it is challenging to assign the regions of density due to their dynamics or structural changes upon binding, making it a much-needed addition to the structural biology toolbox.

## Results

### The cryoENsemble method for Bayesian reweighting with cryo-EM maps

To derive an ensemble of structures, each with a corresponding set of weights that adequately represent the experimental data, we based cryoENsemble on the BioEn method^37,51^ and incorporated a single-particle cryo-EM data framework. The BioEn algorithm uses Bayes’ theorem to define the posterior probability as a function of the statistical weights of each member of the structural ensemble (*w*_*i*_), where *i* is the index of the member, given the experimental data (*D*) and the prior knowledge about the system (*I*)

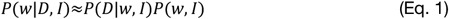

In the context of cryo-EM data, the experimental data points are defined as a set of voxels with a density exceeding a predetermined threshold value, the latter established based on the noise levels present in the data (see Methods). The likelihood function, *P*(*D*|*w, I*) assesses the probability of observing a given set of experimental data (*D*), considering the actual ensemble of structures and their corresponding statistical weights (*w*). The prior probability term, *P*(*w*|*I*), encapsulates the knowledge about the structural ensemble and weights (*w*). This knowledge is typically derived from the molecular dynamics ensemble prior to the incorporation of the experimental data. The prior can be constructed in several ways, but keeping in line with the BioEn methodology, we utilise Kullback-Leibler (KL) divergence (*S*_*KL*_)

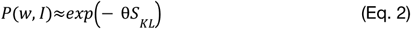

where *S* is defined as 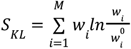 and both reference 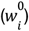 and refined weights (*w*_*i*_) are normalised and positive. Generally, the reference weights of the prior structural ensemble 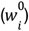 are selected from the uniform distribution, though they can also be set according to populations derived from biased MD simulations (e.g. from metadynamics^52^). An additional hyperparameter, θ, describes our confidence in the initial structural ensemble. A high value of θ indicates high confidence in the MD simulations and generated ensemble, causing the refined weights (*w*_*i*_) to stay close to the initial ones 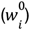. Conversely, a low value of θ, suggests that the initial ensemble may be far from optimal, allowing the weights (*w*_*i*_) to deviate significantly from the starting one 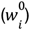. θ is automatically selected during optimisation based on the developed automatic L-curve analysis (see below).

The likelihood function is modelled via a Gaussian distribution^53^

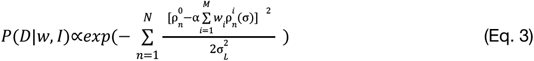

where 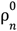 represents the experimental/reference density from the *n-th* voxel of our map, whereas 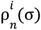 is simulated density from the same voxel generated from the *i-th* model of the structural ensemble with the use of Gaussian functions with the width equal to σ, which is a nuisance parameter (see Methods). This likelihood function contains two additional parameters: the variance of the Gaussian likelihood 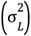, which is equivalent to the experimental error, and the scaling factor (α). We approximate 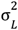 using the variance of noise distribution outside of the molecular system density, while α and σ are estimated simultaneously with the weights during the reweighting. To determine the optimal value of θ, we perform calculations over a range of θ values and use an automatic L-curve analysis with the Kneedle algorithm^54^. This allows the selection of a θ value that yields good agreement with experimental data (low ^2^ χ) and also prevents overfitting (maintaining a small difference from the distribution of the initial weights).

Having defined both the likelihood and prior functions, we can express the log-posterior function, which we will minimise to find the optimal weights, along with nuisance parameters, using the log-weights optimisation as encoded in BioEn

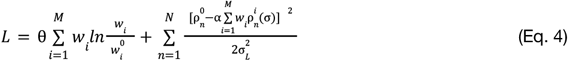

The execution of cryoENsemble calculations yields optimal (non-zero) weights for every structure in our structural ensemble, along with the values of θ, σ and α. A schematic of our methodology is shown in **Fig. 1**.

The method has been validated using two extensive synthetic cryo-EM datasets from the adenylate kinase and ribosomal nascent chain complex (**Fig. 2**, see Methods) and showed that it can accurately reproduce the structural properties of the underlying conformational ensembles from the heterogeneous and noisy cryo-EM data (see Methods).

### Dynamics of the ribosome-bound TF complexed with PDF

Upon binding to the ribosome, TF remains highly dynamic, making it a challenging system for structural studies. We applied the cryoENsemble methodology to the cryo-EM map, which represents the *E. coli* 70S ribosome in complex with PDF, TF and MetAP^38^. PDF and MetAP also bind around the ribosomal exit tunnel and compete for the same binding site localised at uL22-uL32 protein region^47^. MetAP additionally has a secondary binding site, which overlaps with the TF one^38,47^. When TF is bound to the ribosome in the presence of PDF or MetAP, it exhibits reduced dynamics and is, therefore, easier to characterise via cryo-EM.

We exploited this and ran a long all-atom structure-based^55^ MD simulation with TF bound to the surface of the 70S ribosome complexed with PDF (Methods). Despite the fact that the dynamics of TF is restricted in the MD simulations by the presence of the bound PDF, it remains mobile, in particular within the peptidyl-prolyl *cis-trans* isomerase (PPI-ase) domain region (**Supplementary Fig. 1**). We next reweighed the MD trajectory using cryoENsemble and the available cryo-EM map (EMDB: 30611^38^) (**Fig. 3B**) (Methods). Our initial ensemble was already in good agreement with the cryo-EM data, with a correlation coefficient (CC) of 0.68 and Fourier shell correlation (FSC05) of 0.058. Upon reweighting, the agreement improved to CC = 0.71 and FSC05 = 0.091, respectively (**Supplementary Fig. 2**). Moreover, as the reweighting process increased the weights of selected members of the ensemble (**Supplementary Fig. 2**), we identified the best-fit model within the density map, with CC=0.65 (**Fig. 3C**). The improvement of the agreement with the experimental data for the MD ensemble, upon the reweighting, underscores the significance of utilising a structural ensemble instead of a single model when analysing heterogeneous cryo-EM maps. The most significant change in the ensemble composition was evident in our clustering analysis (see Methods). We found that, of the six main clusters encompassing 77% of the total trajectory, only two had an increased population following the reweighting. The remaining four experienced decreased populations, particularly apparent for cluster 1 (**Fig. 3A**), comprising 36% of the MD trajectory.

**Figure 3.**
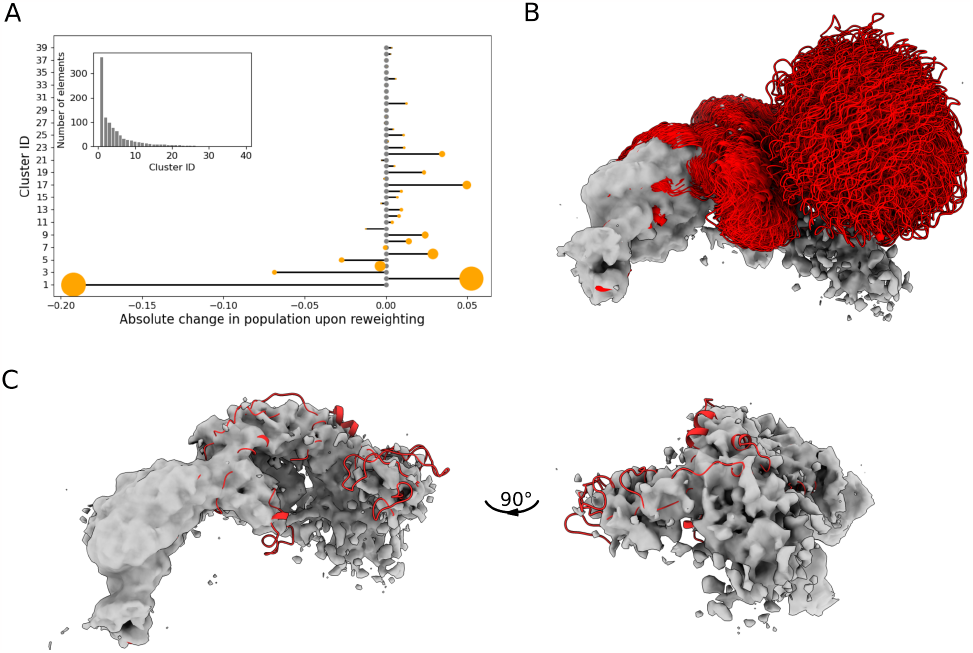
CryoENsemble reweighting of the TF dataset. **(A)** Analysis of the effect of reweighting on the weights of each cluster obtained from the MD simulations. The orange circle size is proportional to the population of the cluster upon reweighting. **(B)** The TF MD ensemble used for reweighting fitted into the cryo-EM map (EMDB: 30611^38^). **(C)** The structural model with the highest weight selected by cryoENsemble (Supplementary Fig. 2) is visualised in two different orientations inside the cryo-EM map.

When we analysed each cluster separately, we noticed that clusters 1 and 2 displayed similar average CC values (0.661±0.017 and 0.659±0.015, respectively), which might have led to equalising their populations via cryoENsemble reweighting and thereby lowering and raising their populations, respectively (**Supplementary Fig. 3**). Both clusters exhibited the highest average value of CC amongst all main clusters. Among the remaining four clusters, cluster 6 has the highest average agreement with the experimental data (CC=0.649±0.019), and its population also increased through reweighting. Interestingly, cluster 3 was observed to experience significant population loss, presenting the lowest CC amongst the main clusters (CC=0.594±0.026) and a visually poor fit into the cryo-EM density (**Supplementary Fig. 3**). Altogether, these findings show that reweighting using cryoENsemble can significantly improve the quality of the MD ensemble and its agreement with the cryo-EM data. Importantly, the reweighting process is not a simple increase of weights for structures with high CC, as we found no correlation between new weights and corresponding CC scores (**Supplementary Fig. 4**) – supporting our observation that an ensemble explains heterogenous cryo-EM data better than a single structure.

Additionally, when we compared the experimental cryo-EM map with the map obtained from the single best-fit model and the map representing the entire reweighted MD ensemble, the reweighted map is more similar to the cryo-EM when visualised at different density thresholds (**Supplementary Fig. 5**). This emphasises not only the necessity to use an ensemble of structures instead of a single structure to capture information from highly heterogeneous cryo-EM maps but also underscores the importance of reweighting.

### Unaccounted cryo-EM density corresponds to a TF-bound methionine aminopeptidase, not TF dynamics

The initial study of TF, MetAP and PDF assembly on the ribosome provided several low-resolution cryo-EM maps of the 70S ribosome in various configurations^47^. Notably, in the cryo-EM map of MetAP-PDF-TF (12.2 Å, EMDB:9778), the MetAP density was unannotated, and an additional density near TF was attributed to the dynamics of TF. A subsequent study obtained a higher resolution cryo-EM map (4.1 Å) of the 70S ribosome with MetAP, PDF and TF with again additional density near TF, but now annotated as a tertiary binding site for MetAP^38^.

Seeking to clarify the nature of this additional density, we took advantage of the unique combination of the MD simulations and cryoENsemble. After accounting for the TF structural ensemble obtained upon reweighting, we observed that there is still an unaccounted density present (**Fig. 4A**), which confirms the suggestion of a tertiary binding site for MetAP^38^. To further validate this observation, we fitted the MetAP structure using a rigid-body procedure, starting with the orientation where the positively charged loops faced the ribosome surface, as indicated by biochemical studies to be a probable ribosome-binding mode^57^, and found a compelling overlap (**Fig. 4, Supplementary Fig. 6**).

**Figure 4.**
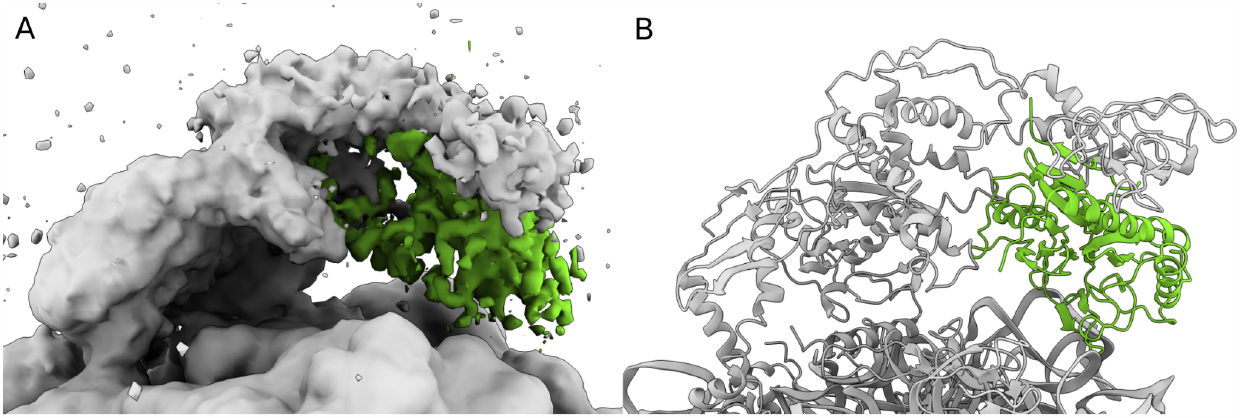
Unaccounted cryo-EM map with corresponding fitted MetAP structure. **(A)** The cryo-EM map (EMDB: 30611^38^) with unaccounted density coloured in green. **(B)** The outcome of fitting the MetAP structure (PDB ID: 1MAT^56^) into the unaccounted density (from **(A)**), presented along the 70S-Trigger factor structure (PDB ID: 7D80^38^).

These findings demonstrate how MD simulations, in combination with cryoENsemble reweighting, can help explain unmodelled and unaccounted for parts of cryo-EM density maps corresponding to dynamic regions of biomolecular complexes.

## Discussion

Characterising complex biological processes through cryo-EM presents many unique challenges, especially for systems that are dynamic or exist in multiple conformational states. We have proposed a method that takes advantage of both the molecular dynamics simulation and Bayesian methodology to yield accurate representations of these systems. This is achieved by reweighting the MD structural ensemble in accordance with cryo-EM data. Notably, unlike most of the existing methods, we do not fit or refine a single structure but adjust the weights of the pre-existing structural ensembles to improve their agreement with the cryo-EM data.

The effectiveness of our approach depends on the quality of the prior structural ensemble since our reweighting strategy by construction does not produce new conformations. In this study, we used all-atom structure-based models^55^ to sample available conformational space efficiently. While structure-based potentials have been previously used to fit structures in the cryo-EM maps via MDFit^58^, we have instead employed them to generate a prior structural ensemble. Despite approximations, structure-based models provide the opportunity to explore the dynamics of large biological complexes, that are inaccessible to more detailed computational approaches and can produce an accurate description of their functional dynamics^59,60^. Coarse-grained simulations, upon initial converting of structures to the all-atom resolution, could be used in a similar manner to generate prior structural ensembles for cryoENsemble, thereby further expanding the accessible system size and complexity.

For more detailed systems, the use of more advanced force fields, such as CHARMM36m^61^ or DES-AMBER^62^, may be necessary to generate more apt initial structural ensembles – potentially even guided by density-driven MD simulations^19^, where lower resolution map or one of the half-maps can be used to restrict sampled conformational space. Moreover, to further increase the capacity to extensively sample the conformational landscape, the structural ensembles derived from MD simulations with enhanced sampling methods can also be used. In this scenario, weights obtained from the reweighted MD simulations would serve to define our initial ensemble. Essentially, any free-energy landscape sampling method, including machine learning, can be used to generate the prior structural ensembles.

CryoENsemble can also be used to improve or validate molecular dynamics simulations. By setting up multiple simulations with different force fields, cryoENsemble can assess which of the force fields most accurately captures the cryo-EM data. Furthermore, this method could be integrated into various force field parameterisation schemes, thereby enabling the utilisation of cryo-EM data^63–65^.

The cryoENsemble approach is particularly suited for complex biological systems featuring convoluted dynamics. These systems often yield cryo-EM maps with high-resolution regions associated with more static components and lower-resolution and ambiguous cryo-EM density describing dynamic elements. Instances of this include nascent chain polypeptides or ribosome auxiliary factors bound to the ribosome. In these cases, the rigid and well-resolved structure of the ribosome contrasts with the low-resolution cryo-EM density of the NC or auxiliary factor. The dynamic character of these components implies the search for a solution in the form of a true structural ensemble rather than a selection of structures, which individually fit into the density or just a single structure^35^.

While our method is computationally efficient, the reweighting time and memory usage can depend on the size of both the cryo-EM map (number of voxels) and the structural ensemble. This can be mitigated by clustering the MD structural ensemble before the reweighting to eliminate highly similar structures, as each requires calculating and storing a density map. The largest structural ensemble we tested comprised 1000 structures, which should suffice to capture the heterogeneity present in the cryo-EM reconstructions for most of the cases. We used a maximum of ∼30,000 voxels from the cryo-EM map; however, one can apply initial down-sampling of the map to reduce the number of voxels for particularly large datasets. This method can also be helpful when working with large structural datasets. An iterative reweighting with a downsampled map can be applied to obtain a minimal set of structures, which can be subsequently reweighted again using the high-resolution map. In a similar fashion, the cryo-EM density can be split into sections with varying resolutions and noise levels or be utilised through separate half-maps. Reweighting the structural ensemble first to the much more refined density and then subsequently reweighting it with a less resolved map region could therefore help mitigate some of the challenges with highly heterogeneous maps.

## Conclusions

We have reported a Bayesian-based approach that enables fitting structural ensembles of various complex biomolecules, including proteins, RNA, DNA, lipids or sugars, into cryo-EM maps to capture both continuous and discrete structural heterogeneity. Through the use of two synthetic datasets and one experimentally-derived cryo-EM map, we have demonstrated that cryoENsemble can generate structural ensembles with averaged density maps closely mirroring the experimental maps, and accurately reproducing the structural properties of the underlying conformational ensembles. We have demonstrated that a fitted structural ensemble captures experimental data better than a single structure in these cases. We have also shown how this method can be applied to analyse unaccounted densities. By enabling the analysis of cryo-EM maps for regions that are more dynamic and therefore have less well-defined density, our method opens up new avenues for structural studies. Additionally, cryoENsemble can be extended to utilise other experimental data within the BioEn or similar framework, making it a potent tool for integrative structural biology.

### Materials and Methods

We validated cryoENsemble using two synthetic datasets. The first one is adenylate kinase (ADK) in both the open and closed conformations, capturing the discrete heterogeneity present in cryo-EM maps. The second system is a ribosome-bound nascent chain of the immunoglobulin-like domain (FLN5-6+31), exemplifying continuous heterogeneity. Characterising ribosome-bound nascent chains using cryo-EM is especially challenging, given that they combine flexible and predominantly unstructured linkers in the exit tunnel with folded or partially folded domains outside of the exit tunnel^66^; the latter only transiently interact with the ribosome^67^.

### Generating the synthetic reference density maps

In our Bayesian framework under typical circumstances, the reference density map would correspond to the experimentally derived cryo-EM map (ρ^*ref*^ *=* ρ^*exp*^) . However, to test our methodology, we utilised synthetic reference density maps. They were either generated based on the crystal structures of the open or closed ADK state (ρ^*ref*^ *=* ρ^*X*−*ray*^) or on randomly selected models from the all-atom MD ensemble of the FLN5-6+31 ribosomal nascent chain 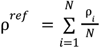, where ρ_i_ is a density map of the i-th model). These synthetic reference density maps were generated using a protocol that mimics *molmap* command from ChimeraX with the bandwidth of the blur kernel σ set at 0.225 × resolution^68^. Maps were produced at three differing resolutions (3, 6, and 10 Å) to explore the influence of resolution on the reweighting process. To further investigate the effect of noise on reweighting, we added different levels of Gaussian noise to the map. The noise had a mean of 0 and a standard deviation based on either 1% or 10% of the map’s maximum density.

### Generating the density maps for the prior structural ensemble

In addition to the synthetic reference density maps (ρ^*ref*^), we generated density maps for every structure from the MD ensemble (ρ^*i*^). If not initially aligned, each structure was aligned to the reference density map using Situs^14^. Following this alignment and using an approach similar to one from the modified *gmconvert* script^32^, density maps were generated with the same voxel size, number of voxels and origin as the reference density map. The process involved positioning a spherical 3D-Gaussian function at each atom position with parameters for the corresponding atom obtained by fitting the electron atomic scattering factors specific to each atom type^32,69^.

### Synthetic density map processing

The generated density maps, both the reference and those from the structural ensemble, were further processed using *mrcfile* python library^70^. From our reweighting dataset, we excluded voxels with negative values and rescaled the remaining ones to a molecular density value of 1, making the different maps easier to compare. Our reweighting methodology operates only on the selected voxels, both from the reference density map and the density maps generated based on the MD ensemble, that have density above the corresponding thresholds. The reference density map threshold is set up to be equal to 3 * σ_*noice*_, where σ_*noice*_ was either 1% or 10% of the maximum density, whereas the threshold for maps generated based on MD was equal to 3*σ_*map*_, where σ_*map*_ is the standard deviation of the synthetic map (**Supplementary Fig. 7**)

### Generation of adenylate kinase synthetic cryo-EM densities

The adenylate kinase is an enzyme that catalyses the phosphoryl group transfer from ATP to AMP. It consists of three domains (CORE, NMP and LID) and undergoes a significant conformational change from open (apo) to closed (holo) conformation upon ligand binding, with RMSD = 7.16Å (**Fig. 2**). Both states have been structurally characterised by X-ray crystallography, with PDB IDs: 1AKE for the closed^49^ and 4AKE^48^ for open conformation. In our study, we generated synthetic density maps based on these X-ray structures, and for the final reference map, we averaged the different populations of open and closed states maps, starting from fully open state conformation and changing the population progressively using 10% intervals until the fully closed conformation was arrived at. In our validation protocol, we operated under the assumption that during the cryo-EM image processing, these states could not be separated into individual 3D reconstructions but were averaged into a single one. We produced 11 averaged reference maps at three different resolutions (3Å, 6Å or 10Å) and with varying levels of Gaussian noise (with a mean of zero and a standard deviation corresponding to 1% or 10% of the maximum ADK density) (**Fig. 5**). In total, we generated 66 synthetic maps for analysis.

**Figure 5.**
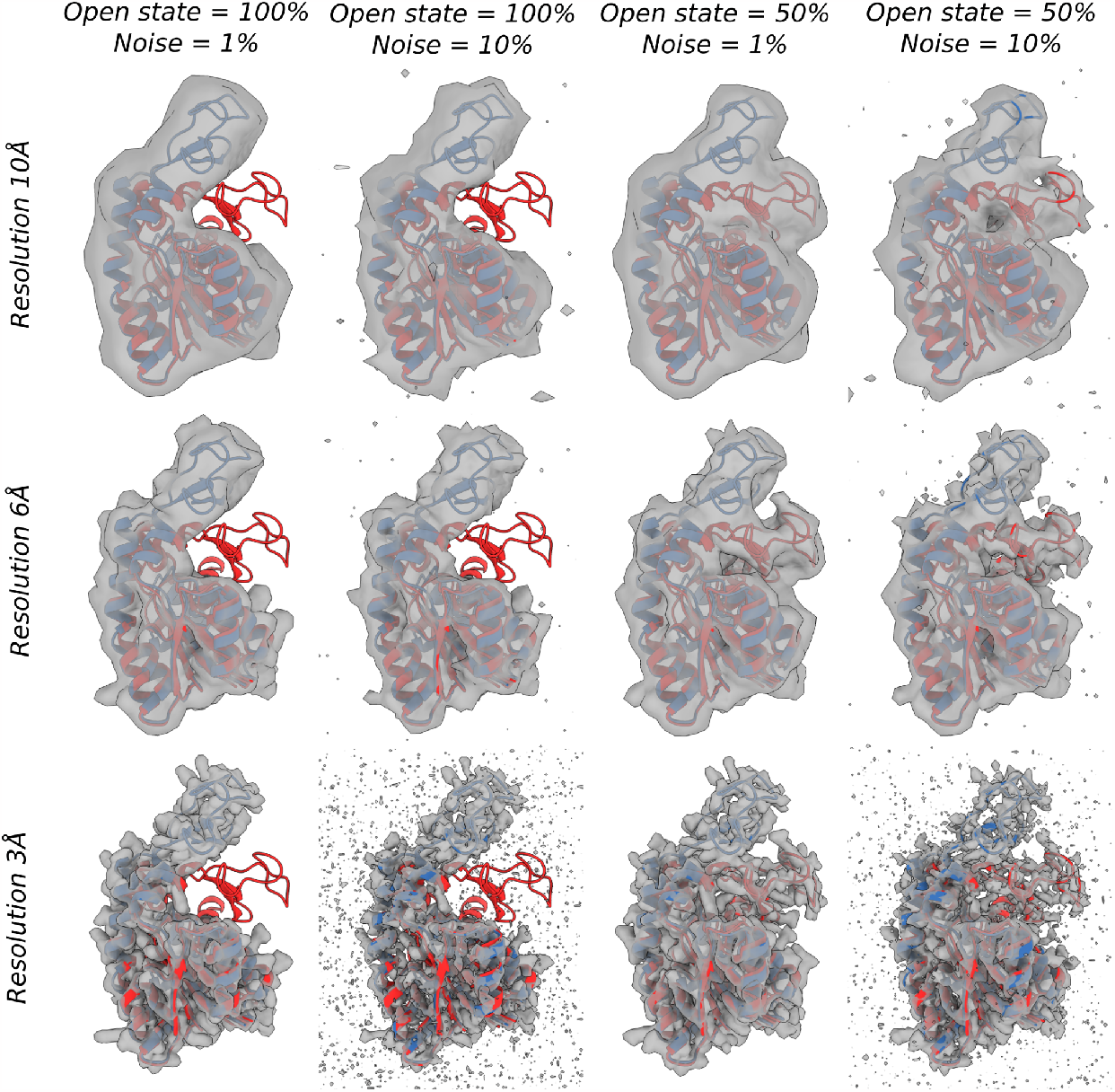
Representative structures and density maps of ADK. ADK X-ray structures in the open (shown in blue) and closed (shown in red) states, along with their generated density maps. The reference density maps were generated based on the varied populations of the open and closed states, different map resolutions, and noise levels. All density maps are depicted at a threshold level equal to three times the standard deviation of the noise distribution.

### Generation of the prior structural ensemble for adenylate kinase

To perform reweighting, we used an MD ensemble of ADK MD, consisting of structures obtained from two short (1.5*10^7^ steps) structure-based all-atom MD simulations with native contacts defined with the Shadow map algorithm^71^ based on the X-ray structures of either the open or closed state. These two ensembles encapsulate the local dynamics around the native state of the apo or holo form. The structure-based models were generated in SMOG 2.0^55^ using all-atoms templates^72^, and the MD simulations were carried out in Gromacs 4.5.7^73^. The combined structural ensemble consists of a total of 100 ADK conformations, with 50 randomly selected from each simulation.

### CryoENsemble reweighting of adenylate kinase dataset

For each ADK dataset, consisting of a structural ensemble and a selected set of voxels from a combination of reference map and simulated map, we ran our cryoENsemble reweighting method. Optimal θ values for each dataset were identified via the L-curve analysis conducted using the Kneedle algorithm, enabling us to compute new optimal weights (see SI). Initially, we assessed the effectiveness of our methodology in reproducing the reference population of the open state used to generate the reference maps (**Fig. 6**). Our findings indicate consistency across all density maps, which decreases with both a reduction in resolution (10 Å) and an increase in the noise level (10%) (**Fig. 6**). Interestingly, the consistency was lower for reference maps with a very small (<=0.2) or very large (>=0.8) population of the open state. This disparity is likely due to our prior structural ensemble equally representing the open and closed states, a significant deviation from these reference maps. This observation underscores the potential of our method to prevent overfitting.

**Figure 6.**
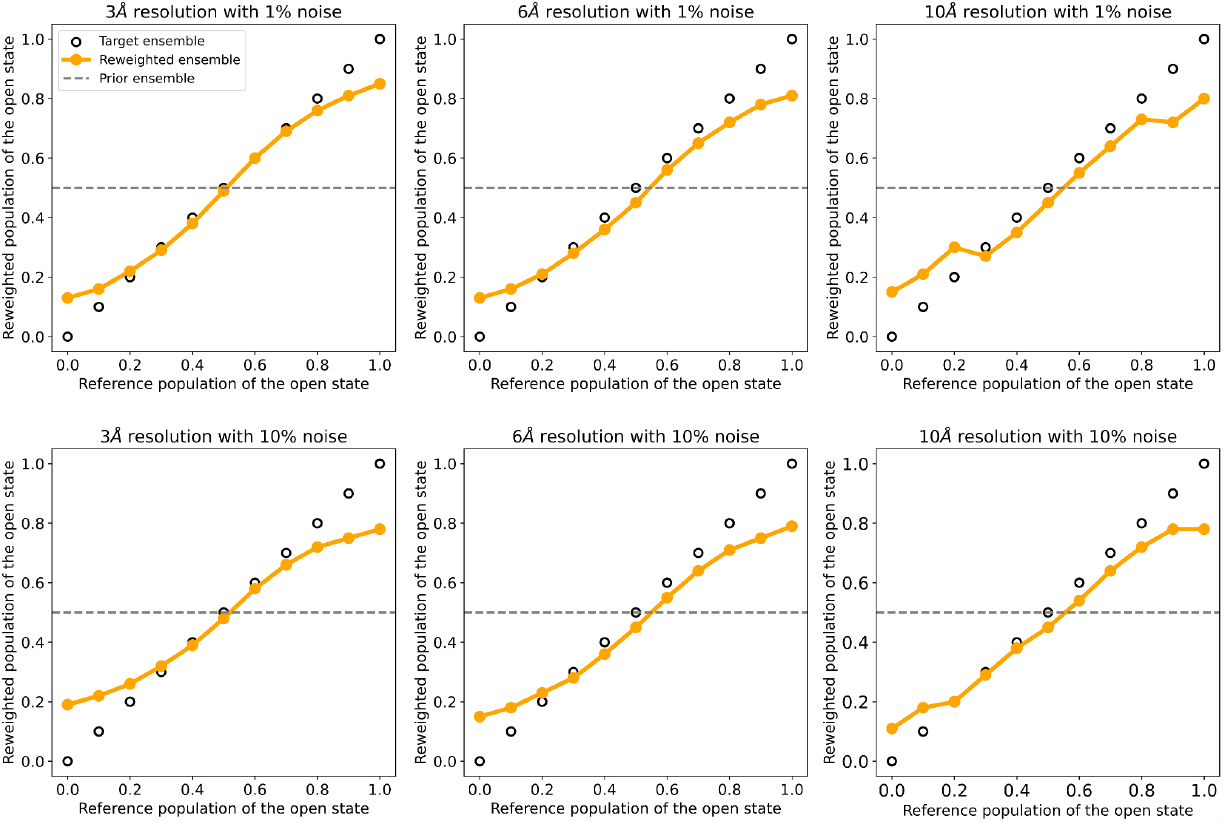
Open state populations calculated from cryoENsemble reweighting of ADK dataset. The open state populations obtained after structural ensemble reweighting for each ADK dataset are shown in orange along the target values (circle). The datasets varied in resolution, noise level, and reference populations of the open state.

Following these calculations, we generated a reweighted and averaged cryo-EM map for each dataset and compared it with the reference density map to evaluate the impact of the reweighting (**Fig. 7**). For the comparison, we applied three different metrics as recommended by the 2019 Cryo-EM Model Challenge^74^. Scores were used based on the correlation coefficient (CC), Fourier shell correlation (FSC05) and Segment Based Manders’ Overlap Coefficient (SMOC)^75^ as they represent the three main clusters of the scoring methods and therefore capture various aspects of similarity between maps. The correlation coefficient calculated between the reference maps and posterior densities revealed that the effect of reweighting is evident across all open state populations in each ADK dataset (**Fig. 7**). The correlation coefficient for medium and low-resolution maps reaches values of up to 0.9 or 0.8 for low (1%) and high (10%) noise levels, respectively. Higher resolution maps (3 Å), which contain more detail, present a more significant challenge in reweighting the structural ensemble to achieve high correlation coefficients, in particular when high noise levels (10%) are present in the data (**Fig. 7**). However, reweighting consistently yields higher correlation coefficients than those obtained with the prior weights (for 3 Å with high noise levels (10%) on average CC=0.597 and CC=0.558 for posterior and prior weights, respectively). We also compared our reweighting results with the correlation coefficients derived from maps generated based on the best single structure fit. In the majority of the cases, the entire ensemble obtained after reweighting provides a more accurate representation of the map than any individual structure (**Fig. 7**). Interestingly, when lowering the map resolution, e.g. from 6 to 10 Å, the quality of a single structure fit increases and in the cases of either entirely open or closed state it is higher than the CC of the prior ensemble, and can even equal the value of the reweighted ensemble. The single structure CC deteriorates when maps are close to an equal mixture of open and closed states. These observations show that cryoENsemble not only can provide a reweighted structural ensemble but also inform on when a single structure may be sufficient to describe a cryoEM map satisfactorily.

**Figure 7.**
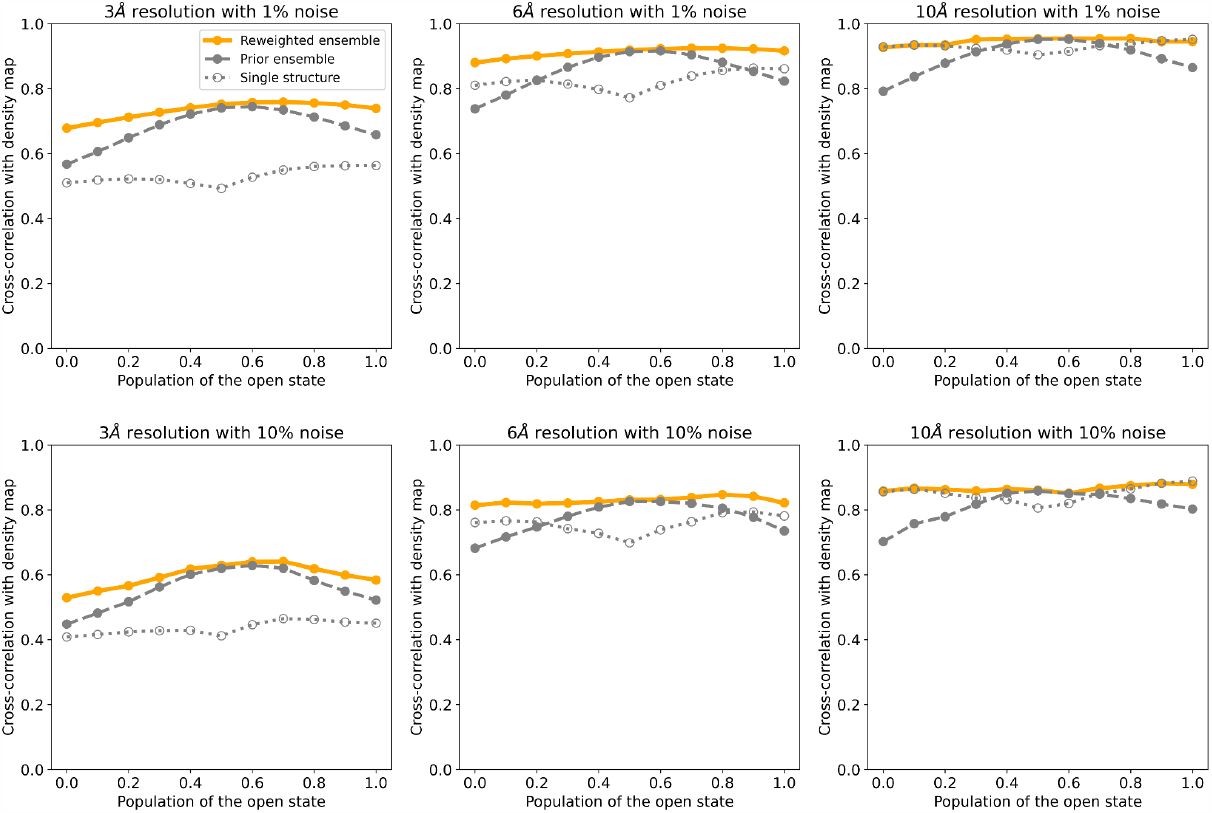
Correlations between reference maps and posterior maps upon cryoENsemble reweighting of ADK dataset. Correlation coefficients calculated between the reference maps and the maps generated from the structural ensemble upon reweighting for each ADK dataset. The datasets varied in resolution, noise level, and reference populations of the open state.

Using a score based on the model-map FSC curve read at FSC=0.5 (FSC05)^76^ (another global score which, unlike CC, increases with the higher resolution of the density map of the target ^74^) on the ADK dataset also showed an improvement upon the cryoENsemble reweighting (**Supplementary Fig. 8**). As a final validation score, we used SMOC that captures the local similarity between the reference map and the fitted model. We calculated the average SMOC score across all residues and models from the MD ensemble and observed that it improved after reweighting, in particular for the entirely open and closed states (**Supplementary Fig. 9**). Altogether, for both global and local metrics, we see a clear effect of the reweighting on the quality of the ADK structural ensemble. We subsequently analysed how the weights of each model from the structural ensemble were updated during the reweighting. We found that cryoENsemble shifted the weights from the uniform prior distribution to correctly capture the reference map open/closed state population (**Supplementary Fig. 10-15**). Finally, a visual comparison of the prior and posterior average density maps alongside the reference maps shows the impact of the reweighting, especially pronounced for the open state maps where posterior maps combine only the open state conformations (**Supplementary Fig. 16-18**).

Overall, we have demonstrated the effectiveness of cryoENsemble in characterising discrete heterogeneity in cryo-EM maps. We derived weights that can generate a map in good agreement with the experimental data (**Supplementary Fig. 16-18**) and are able to describe the correct populations of each state (**Fig. 6**). Our reweighted structural ensemble better explains the experimental data, using both global (**Fig. 6, Supplementary Fig. 9**) and local metrics (**Supplementary Fig. 9**), than the starting ensemble. Furthermore, our method can also suggest when a single structure fitted into the reference map is insufficient, highlighting the necessity of using a structural ensemble for heterogeneous cryo-EM map fitting.

### Generation of the prior structural ensemble and synthetic cryo-EM densities for the FLN5-6+31 RNCs

The FLN5-6 ribosome nascent chain complex encompasses the immunoglobulin-like domain (FLN5 protein), captured during its biosynthesis on the bacterial 70S ribosome. The FLN5-6 nascent chain sequence also consists of the 31 amino-acid linker comprising the fragment of the subsequent filamin domain (FLN6) and the SecM stalling sequence^77^. The FLN5 is the fifth filamin domain (residues 646-750) of the *Dictyostelium discoideum* gelation factor, and its co-translational folding has been extensively studied through a combination of experimental and computational techniques^66,67,78,79^. For our study, we generated a starting ensemble by randomly selecting 100 conformations of the NC from the FLN5-6 structural ensemble obtained from the previous all-atom structure-based MD simulation ^50^ (**Fig. 2** and **Fig. 8**). This ensemble exhibits significant structural heterogeneity, with RMSD values up to 28Å (**Supplementary Fig. 19**), reflecting the dynamic nature of the RNCs. To obtain the reference density maps, we randomly selected ten structures from this starting ensemble, generated an average density map and repeated this procedure 100 times with Gaussian noise, corresponding to either 1% or 10% of the main density added (**Fig. 8**). This system enables us to evaluate our methodology in the case of continuous heterogeneity present in the cryo-EM maps.

**Figure 8.**
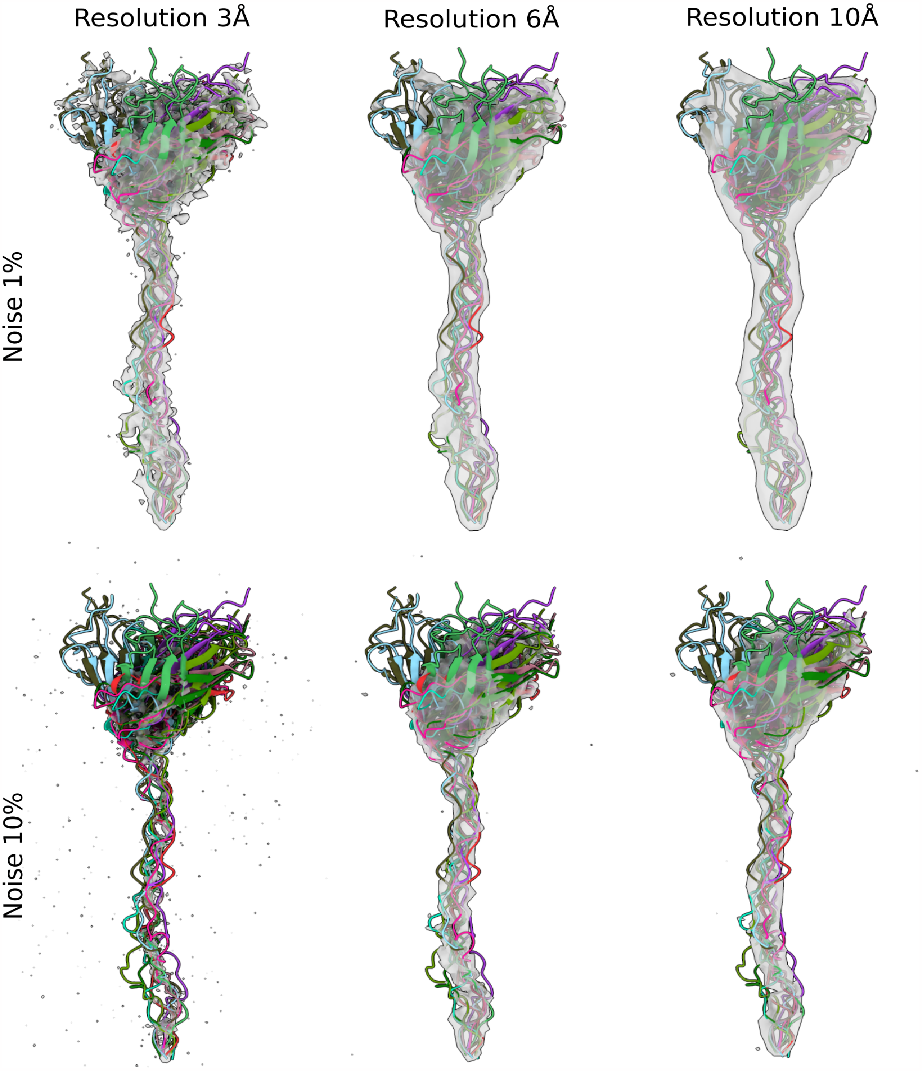
Representative structures and densities of the FLN5-6 nascent chains. Examples of ten random FLN5-6 nascent chain structures chosen from the MD ensemble (**Fig. 1B**) used to generate the reference density map at different resolutions (3, 6, and 10 Å) and noise levels (1% or 10%). Each structure is depicted in a different colour and combines the FLN5-6 nascent chain that is composed of N-terminal folded FLN5 followed by 31 amino acids of the subsequent FLN6 domain and a C-terminal SecM stalling sequence that is covalently attached to the tRNA at peptidyl transferase centre of 70S ribosome. All maps are depicted at a level equal to three times the standard deviation of the noise distribution.

### Reweighting the structural ensemble of the ribosome-bound nascent chain of FLN5-6

We applied cryoENsemble protocol to the prior structural ensemble and optimised the log-posterior function (Eq. 4), using the combined data from the reference map and maps generated from the structural ensemble.

We used the entire structural ensemble (100 models) for the reweighting, including the ten conformations used to generate the reference density maps. Average density maps were generated based on the prior and posterior weights, and their correlation coefficients with the reference density maps were calculated (**Fig. 9**). The prior ensemble, despite its significant structural heterogeneity, already displayed a good agreement with the reference density maps with average CC varying between 0.8 (3 Å maps) and 0.95 (10 Å maps) for 1% noise and from 0.47 (3 Å maps) to 0.84 (10 Å maps) for 10% noise (**Fig. 9**). After reweighting, the correlation increased in all cases, reaching a value close to 1.0 for low noise levels (1%) and 0.9 for high noise levels (10%), with the only exception of the 3 Å maps, which, as in the ADK case, present a more significant challenge in reweighting the structural ensemble, in particular at the higher noise levels (10%) where CC reached 0.54 vs 0.47 with the prior weights. This difficulty is further apparent upon examining the extent of density in this highly noisy system (**Fig. 8**). A comparison with maps generated based on a single structure shows that, in contrast to the ADK system, a single structure cannot represent the dynamic heterogeneity present in the nascent chain cryo-EM maps for any of the systems we tested (**Fig. 9**). We also evaluated the reweighted ensemble using FSC05 and SMOC metrics (**Supplementary Fig. 20 and 21**), finding that the reweighting improved the agreement with experimental data both globally and locally in all cases. Additionally, in order to assess the structural similarity between the obtained reweighted ensemble and the ten structures used to generate the reference map, we used the Jensen-Shannon (JS) divergence. We found significantly closer matching values upon reweighting. For the prior ensemble, the JS divergence was equal to 0.112±0.042 whereas for the posterior ensemble, it varied in 1% noise maps from 0.039±0.016 (3 Å) to 0.065±0.024 (10 Å) and in 10% noise maps from 0.058±0.02 (3 Å) to 0.068±0.026 (10 Å) (**Supplementary Fig. 22**).

**Figure 9.**
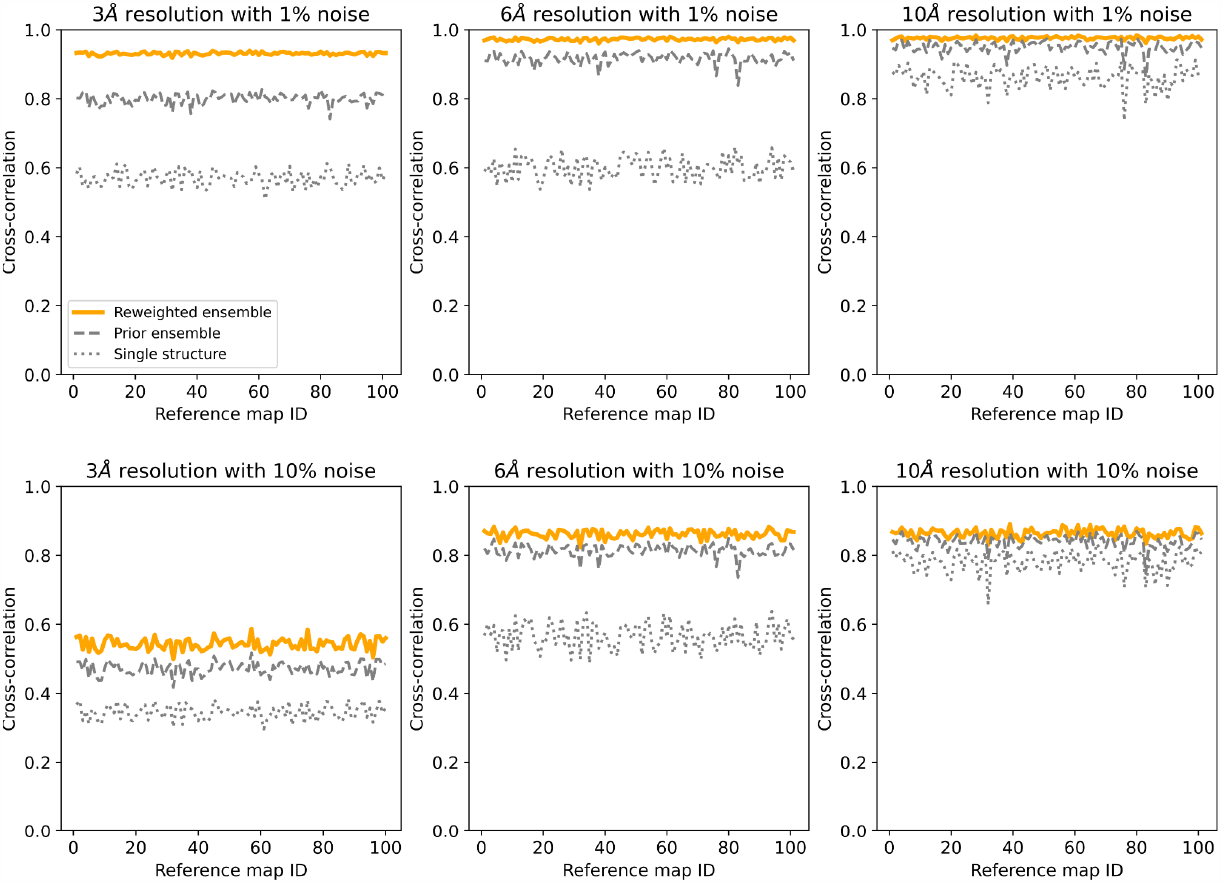
Correlations between the reference maps and posterior maps upon cryoENsemble reweighting of the FLN5-6 nascent chain dataset. Correlation coefficients calculated between the FLN5-6 nascent chain reference density maps and maps obtained before and after the reweighting, as well as the maps derived from the best single structure fitted into the reference density map. The 100 reference density maps (at resolutions of 3, 6, and 10 Å, and with noise levels of 1% and 10%) were generated based on ten randomly selected structures from the MD ensemble.

The high initial correlation between the prior ensemble and reference map can result in relatively minor changes to the correlation coefficients after reweighting (**Fig. 9**). However, we observed significant shifts from the uniform distribution of the weights of the prior structural ensembles due to reweighting (**Fig. 10**). The weights of the ten models used for reference map generation (circled in **Fig. 10**) are substantially higher than those of the remaining structures (e.g. 5.6% vs 0.5% on average for 3 Å maps with 1% noise), a trend not significantly affected by the resolution of the density map or its noise level. This observation highlights the sensitivity of our method, which became particularly apparent when we analysed the entire dataset to determine how many of the ten models used to generate the reference map received the highest weight after the reweighting (**Supplementary Fig. 23**). For high- and medium-resolution maps (3 and 6 Å), our method assigned the highest weights to the correct models in all datasets. While lower-resolution maps (10 Å) posed greater challenges, over half of the reference models were correctly identified to receive the top ten highest weights.

**Figure 10.**
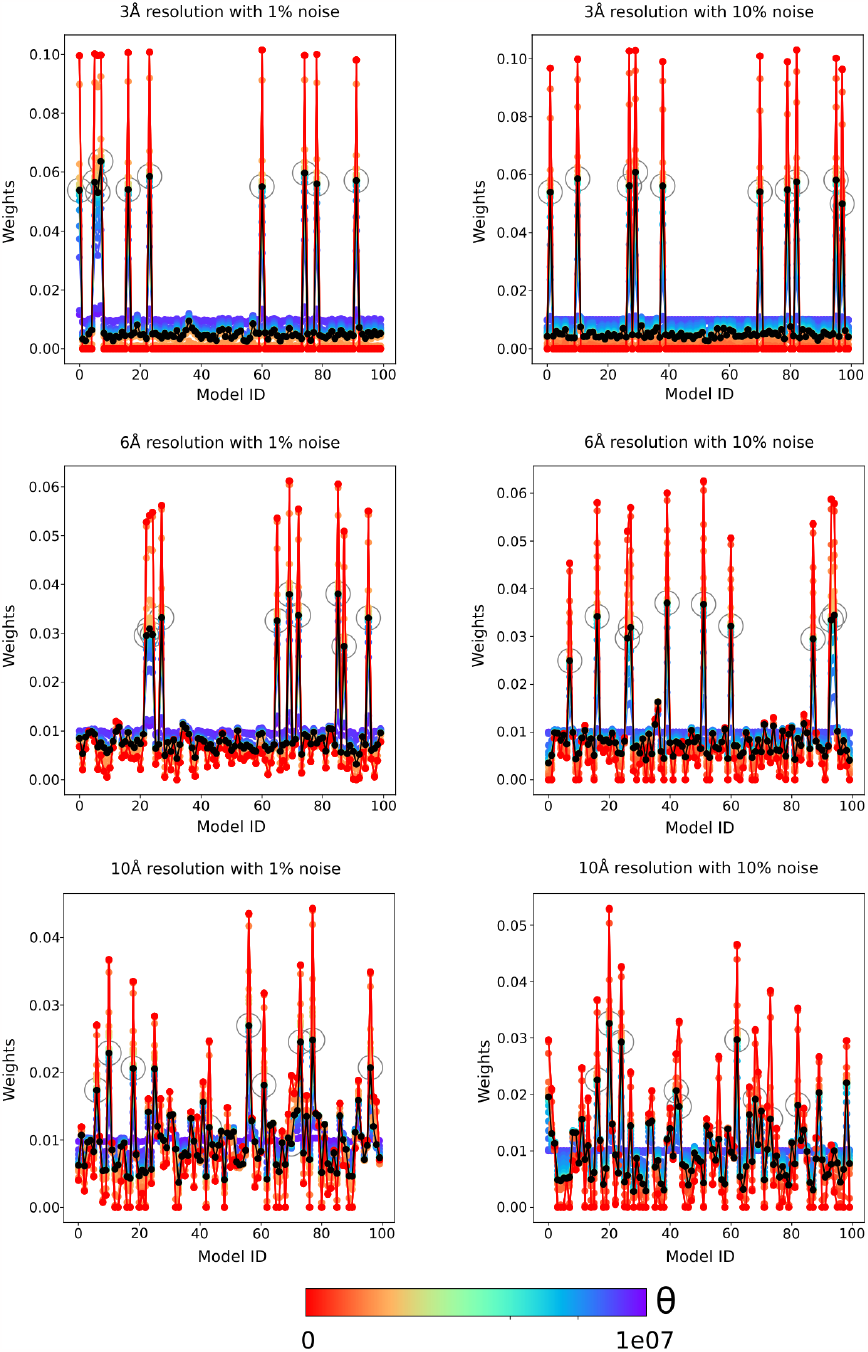
Weights obtained upon cryoENsemble reweighting of the FLN5-6 nascent chain dataset. Examples of the reweighting process for FLN5-6 nascent chain based on the reference map (at resolutions of 3, 6, and 10 Å, and with noise levels of 1% and 10%). Weights are calculated with different theta (θ) values ranging from 0 to 10^7^, and with black lines, we depict optimal weights selected based on the L-curve analysis. Additionally, weights corresponding to the ten models used to generate the reference map are circled.

### Preparation of the trigger factor cryo-EM map for reweighting

For the final system, we used an experimentally derived cryo-EM map capturing the dynamics of the ribosome-associated chaperone (trigger factor) bound to the ribosome in the presence of the peptide deformylase and excess of methionine aminopeptidase (**Fig. 2**)^38^. The obtained cryo-EM map had a clear density for the 70S ribosome, TF and PDF, which enabled authors to fit and refine models. However, the presence of the incomplete MetAP density suggested a novel tertiary binding site but did not allow for modelling the bound state. To create a reference map for the reweighting process, we used ChimeraX to select only the density that corresponds to either the Trigger Factor (TF) or the unmodeled MetAP (**Fig. 2**), subsequently saving it as a smaller, cropped map. This map was then normalised using the same methodology that we previously outlined for the synthetic reference map. We used the map threshold suggested by the authors (0.005) to select significant voxels for the reweighting.

### Generation of the Prior Structural Ensemble for the TF System

Using the available structure of the 70S ribosome from *E*.*coli* with bound TF and PDF (from PDB ID: 7D80^38^), we prepared a starting structure for the MD simulation that encompassed the surface of the 70S ribosome around the ribosomal exit tunnel and bound both TF and PDF (**Supplementary Fig. 24**). We used an all-atom structure-based model generated with SMOG 2.4.4^55,80^ with bond lengths and angles based on the AMBER03 force field^81,82^. Native contacts that are used in structure-based potential were defined based on TF cryo-EM structure with the use of the Shadow Map^71^. For the structure-based MD simulations set up in SMOG, reduced units were applied with length, time, mass and energy scale all set to 1, except for the Boltzmann constant, which is k_B_ = 0.00831451 (kJmol^−1^K^−1^, default in GROMACS). Simulations were performed for 5*10^8^ steps in GROMACS 2021.2^83^ in NVT ensemble at a reduced temperature of 0.5 (60 in GROMACS units), which is slightly below the temperature for this model to capture physiological conditions (0.582 reduced unit^81^). The constant temperature was maintained via the Langevin Dynamics protocol. Taking advantage of a recent comparison of diffusion coefficients in the SMOG model and an all-atom explicit-solvent model^84^, we estimated the effective simulated time to be in a range of hundreds of microseconds. During simulations, we kept the atoms of the ribosome surface frozen. We sampled the trajectory every 5*10^5^ steps generating 1000 structures, and clustered them based on the RMSD using the *gmx cluster* method from GROMACS (**Fig. 3**). Obtained structural ensemble, we used as a prior during the reweighting process carried out in cryoENsemble.

### Fitting of the MetAP

To isolate the MetAP cryo-EM density, we utilized the ChimeraX^68^ command ‘volume subtract’ to create a difference map between the original (EMDB: 30611^38^) and the posterior map derived from cryoENsemble reweighting of the TF MD trajectory. The *E*.*coli* methionine aminopeptidase structure (PDB ID: 1MAT^56^) was fitted into the obtained density using ChimeraX, orienting positively charged loops towards the ribosome, in accordance with previous studies^57^. For subsequent rigid-body fitting, we utilized the ‘Fit in Map’ command, setting the simulated map resolution to 8Å.

## Code Availability

The source code of CryoENsemble, accompanied by a basic tutorial, is freely available on GitHub at: https://github.com/dydymos/cryoBioEN.

## Acknowledgements

J.C. is supported by the Wellcome Trust (Investigator Awards 097806/Z/11/Z & 206409/Z/17/ Z). This project made use of time on HPC granted via the UK High-End Computing Consortium for Biomolecular Simulation, HECBioSim (http://hecbiosim.ac.uk), supported by EPSRC (grants no. EP/R029407/1 and EP/X035603/1). We also acknowledge the EuroHPC Joint Undertaking for awarding this project access to the EuroHPC supercomputer LUMI, hosted by CSC (Finland) and the LUMI consortium through a EuroHPC Regular Access call and the Baskerville Tier 2 HPC service (https://www.baskerville.ac.uk/). Baskerville was funded by the EPSRC and UKRI through the World Class Labs scheme (EP/T022221/1) and the Digital Research Infrastructure programme (EP/W032244/1) and is operated by Advanced Research Computing at the University of Birmingham.

## Author contributions

T.W., J.O.S., A.M., L.D.C., M.V. and J.C. designed the project. T.W. performed the research. T.W., J.O.S., A.M., L.D.C., M.V. and J.C. prepared the manuscript.

