## Supplementary Material for "CryoENsemble - a Bayesian approach for reweighting biomolecular structural ensembles using heterogeneous cryo-EM maps"

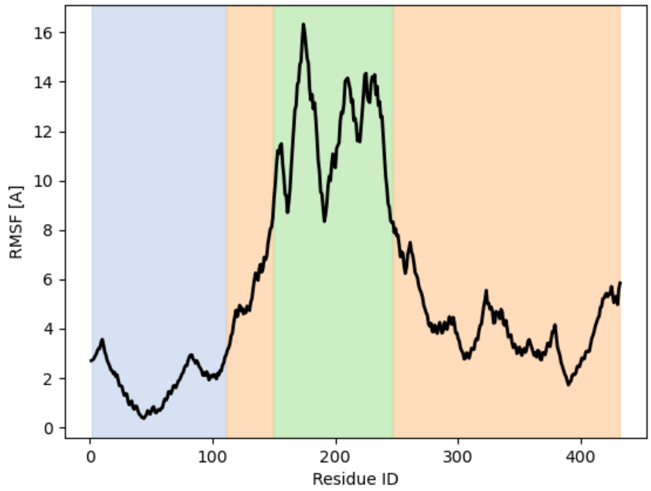


**Supplementary Figure 1**

The root-mean-square fluctuation (RMSF) calculated from a long all-atom MD simulation of the TF bound to the ribosome. Regions corresponding to each domain of the TF are marked with colours: the ribosome binding domain in blue, the peptidyl-prolyl *cis-trans* isomerase domain in green and the subtract binding domain in orange.

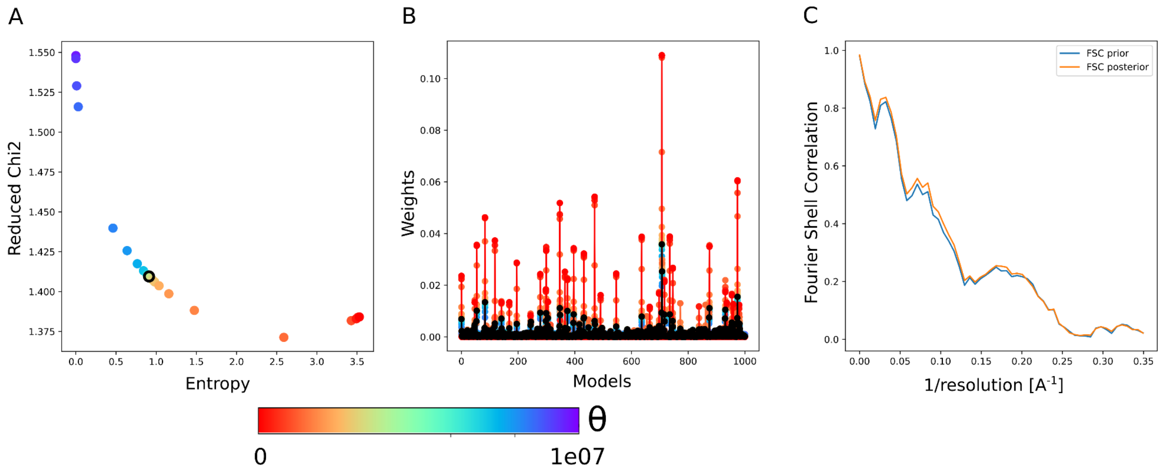


**Supplementary Figure 2**

Results of the trigger factor dataset reweighting. A. The L-curve analysis to select the optimal Θ parameter (in black). B. The reweighted weights for each model from the MD ensemble plotted for different Θ values, in black weights for optimal Θ. C. The Fourier Shell Correlation between the reference experimental cryo-EM map and the average map generated from the MD ensemble, before and after the reweighting.

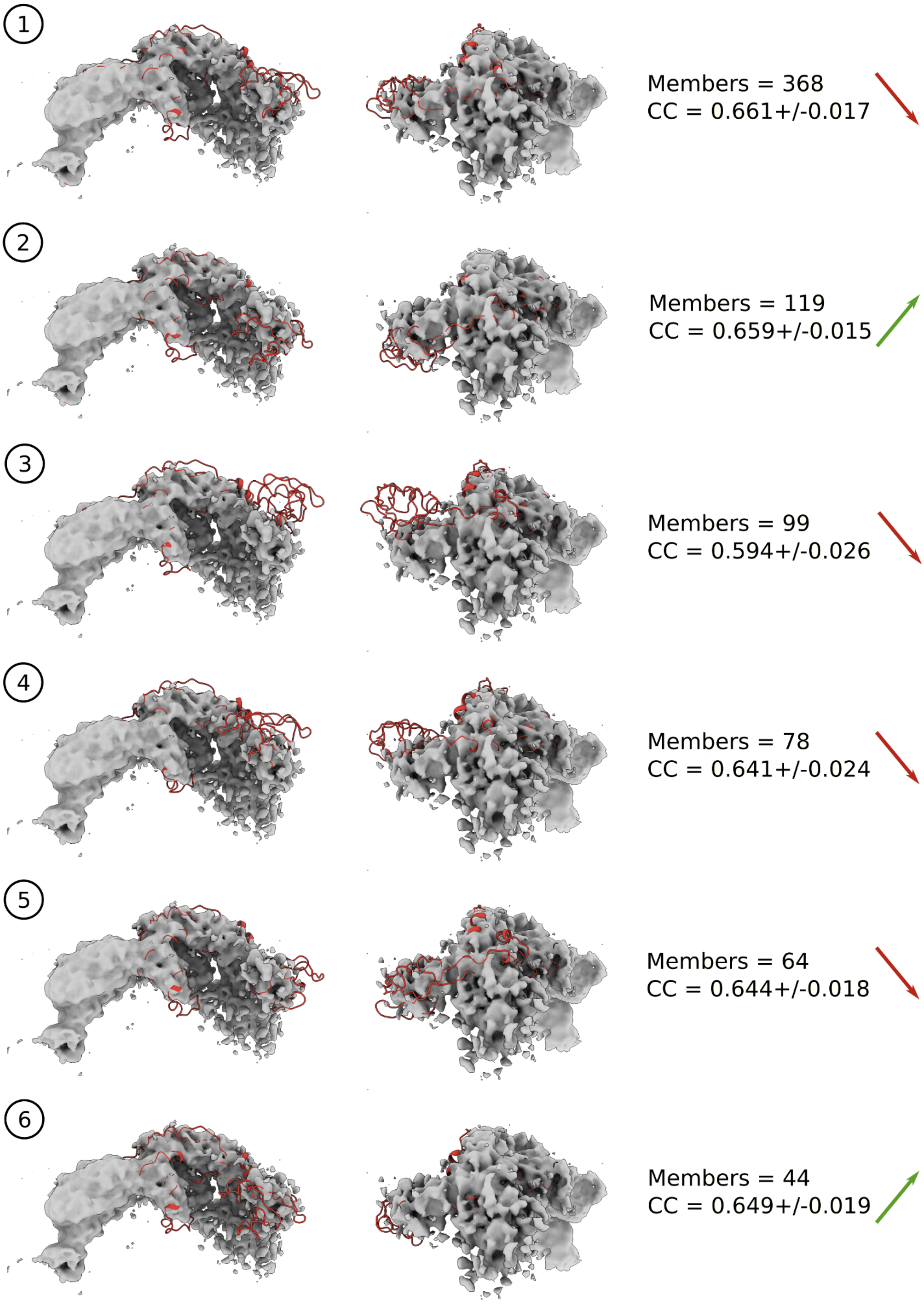


**Supplementary Figure 3**

Experimental cryo-EM density map shown with central structures from the six main clusters obtained from the MD trajectory. Alongside the size, we present the average correlation coefficient calculated for each cluster. The arrows present the change in the population of each cluster upon reweighting, with red indicating a decrease and green increase in the populations.

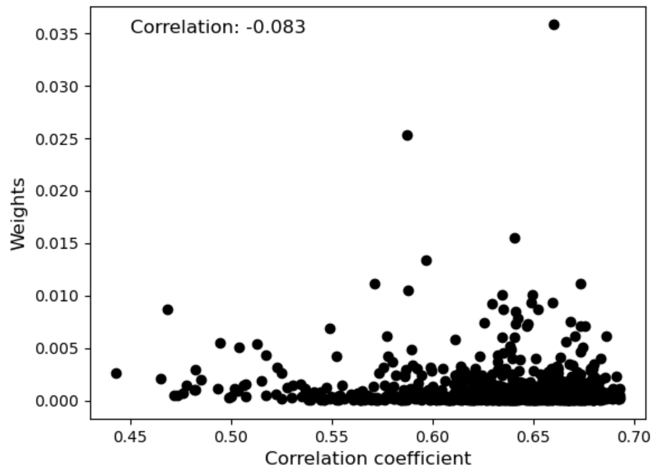


**Supplementary Figure 4**

Correlation between the weights of each model from the trigger factor MD ensemble upon reweighting and its level of agreement with the cryo-EM density expressed by correlation coefficient (CC).

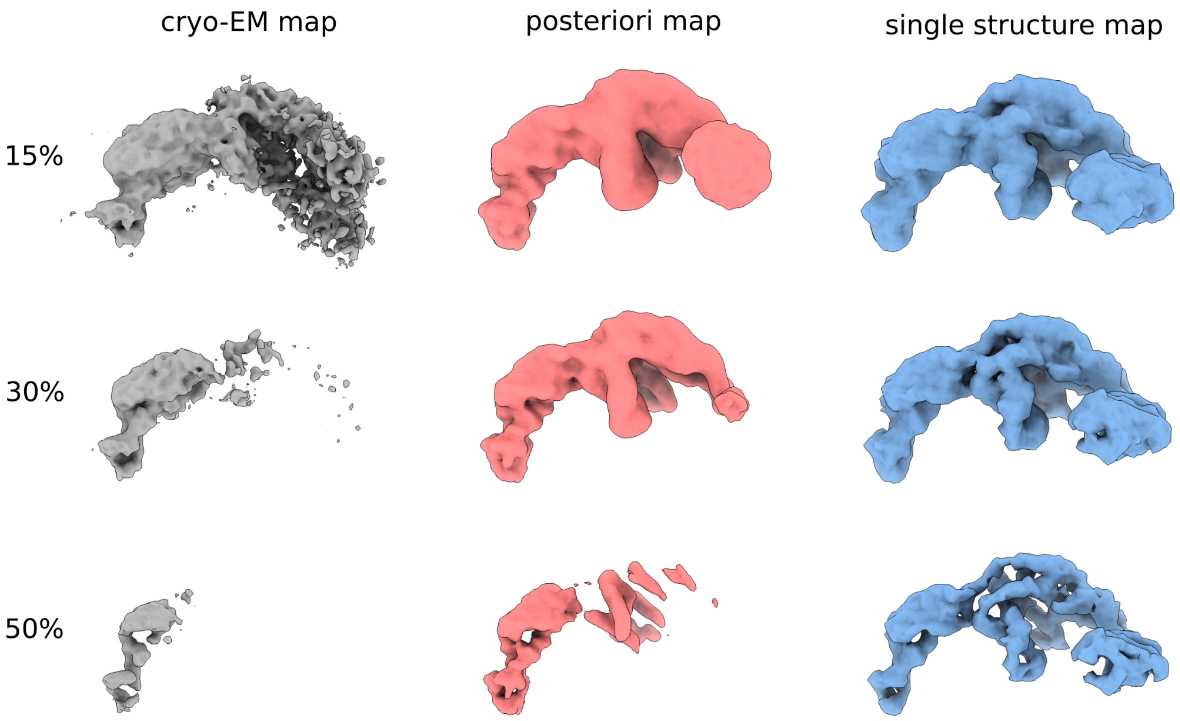


**Supplementary Figure 5**

The trigger factor cryo-EM map (in grey) presented along the calculated map for the posterior ensemble (in light red) and single best-fit structure (in light blue) at different levels of the map threshold. Each map was normalised, to enable comparison, and the selected threshold corresponded to the % of the maximum map value.

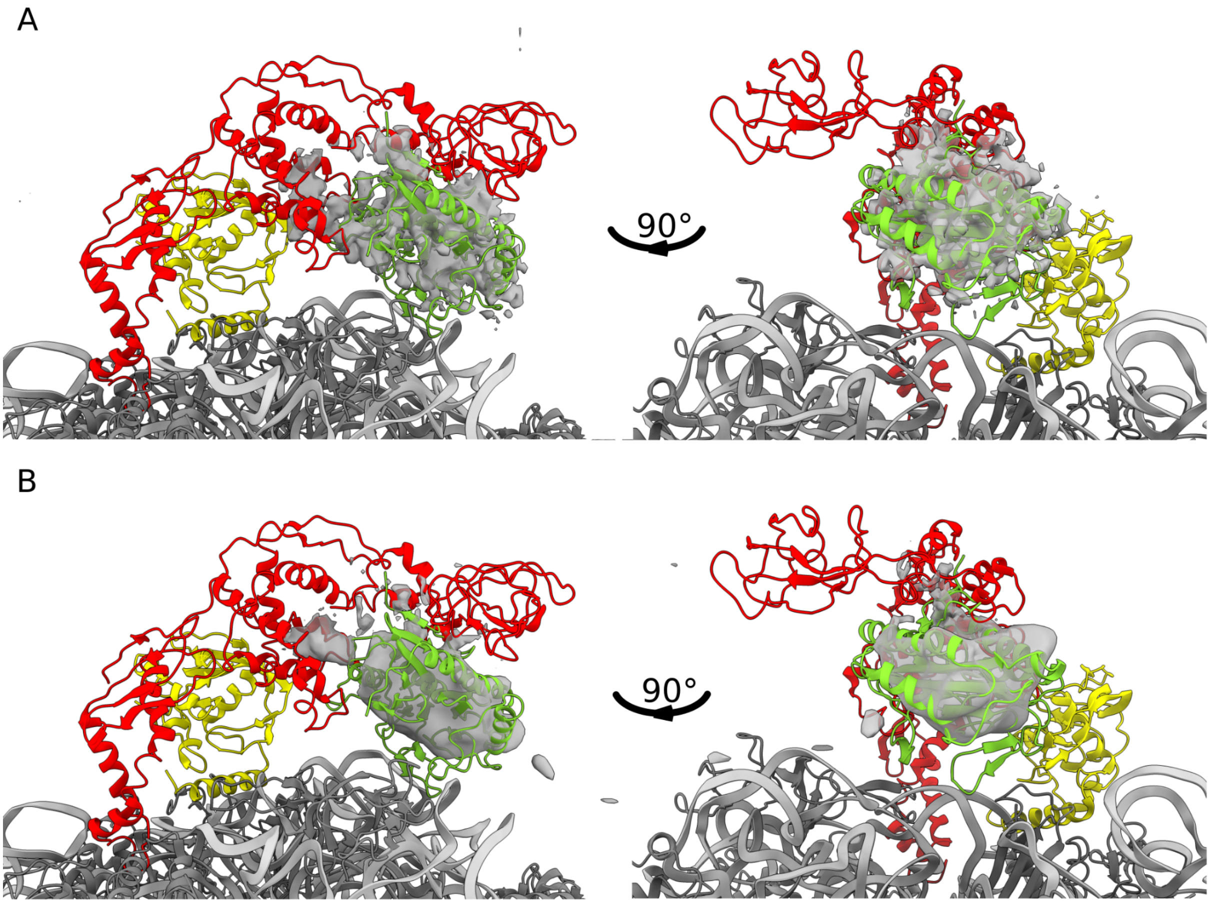


**Supplementary Figure 6**

X-ray MetAP structure (in green) fitted into the unanccounted cryo-EM density (in grey) from (A) EMDB:3061 and (B) EMDB:9778 maps. Additionally we present 70S ribosome in grey (ribosomal protein) and silver (RNA) allong side bound trigger factor in red and PDF in yellow.

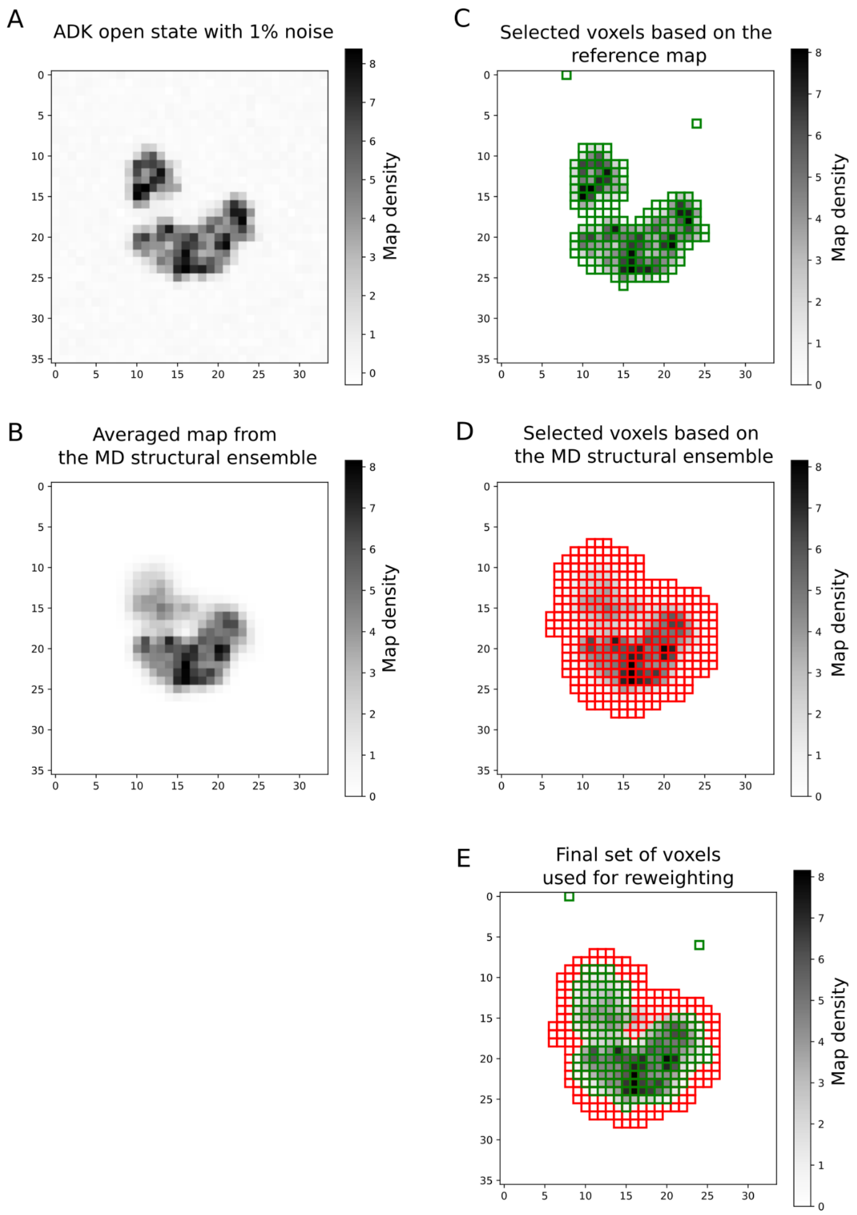


**Supplementary Figure 7**

The 2D cross-sections of the cryo-EM densities from the ADK open state with 1% noise level **(A)** and the averaged map from the corresponding MD structural ensemble **(B)**. **C.** The cross-section from (**A**) with selected (in green) voxels that are above the noise level and will be used for reweighting. **D.** The cross-section from (**B**) with selected (in red) voxels that are above the threshold level and will be used for reweighting. **E.** The cross-section from (**B**) with a selected final set of voxels used for reweighting that comprise ones selected based on cryo-EM reference density as well as from the MD ensemble.

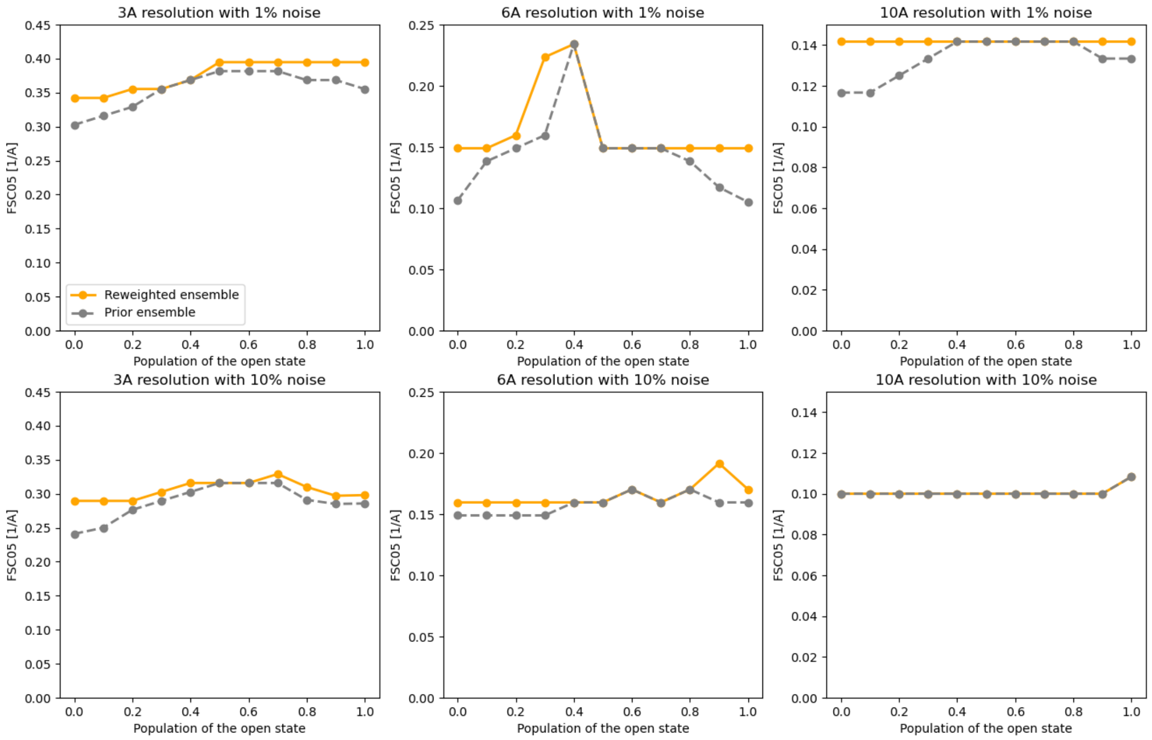


**Supplementary Figure 8**

The FSC05 calculated between the reference map and the maps generated from the structural ensemble upon reweighting for each ADK dataset. The datasets varied in resolution, noise level, and reference populations of the open state. For the 6 Å and 10 Å maps with 1% noise level, we used FSC05 calculated at FSC=0.8 due to too high correlation values.

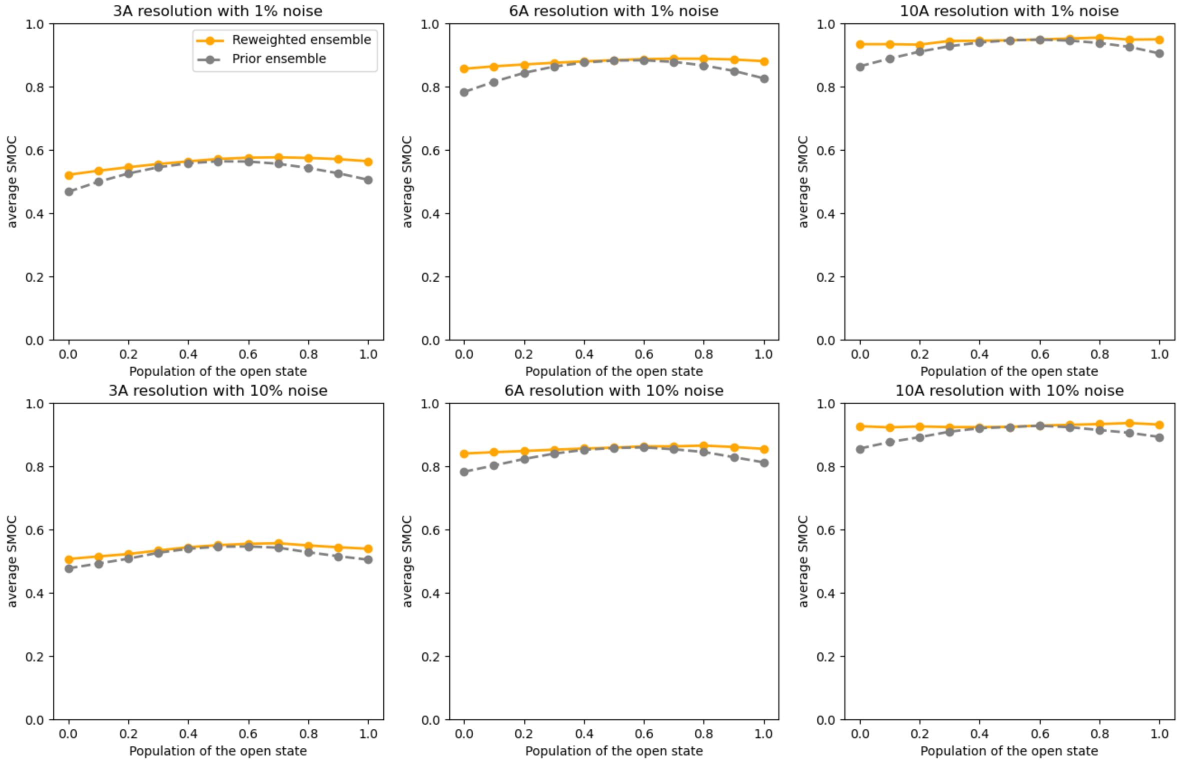


**Supplementary Figure 9**

The average SMOC score calculated between the reference map and the map generated from the structural ensemble upon reweighting for each ADK dataset. The datasets varied in resolution, noise level, and reference populations of the open state.

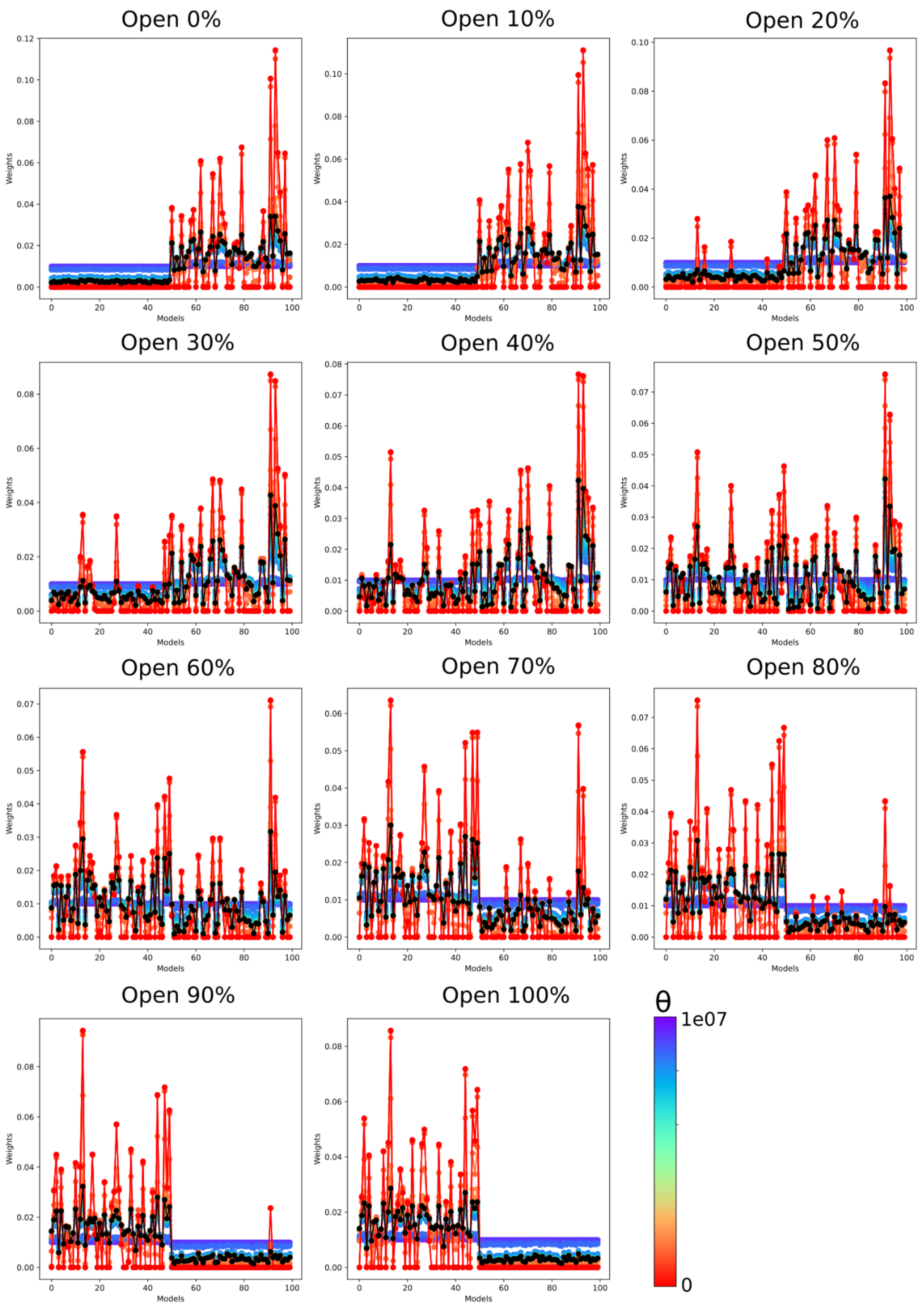


**Supplementary Figure 10**

The weights assigned to each model from the structural ensemble after the reweighting with the use of a 3 Å reference map with the different populations of the open state and a 10% noise level. The first 50 models represent the open conformation, while the last 50 models represent the closed conformation. Weights are presented for various θ values, with the optimal θ value shown in black.

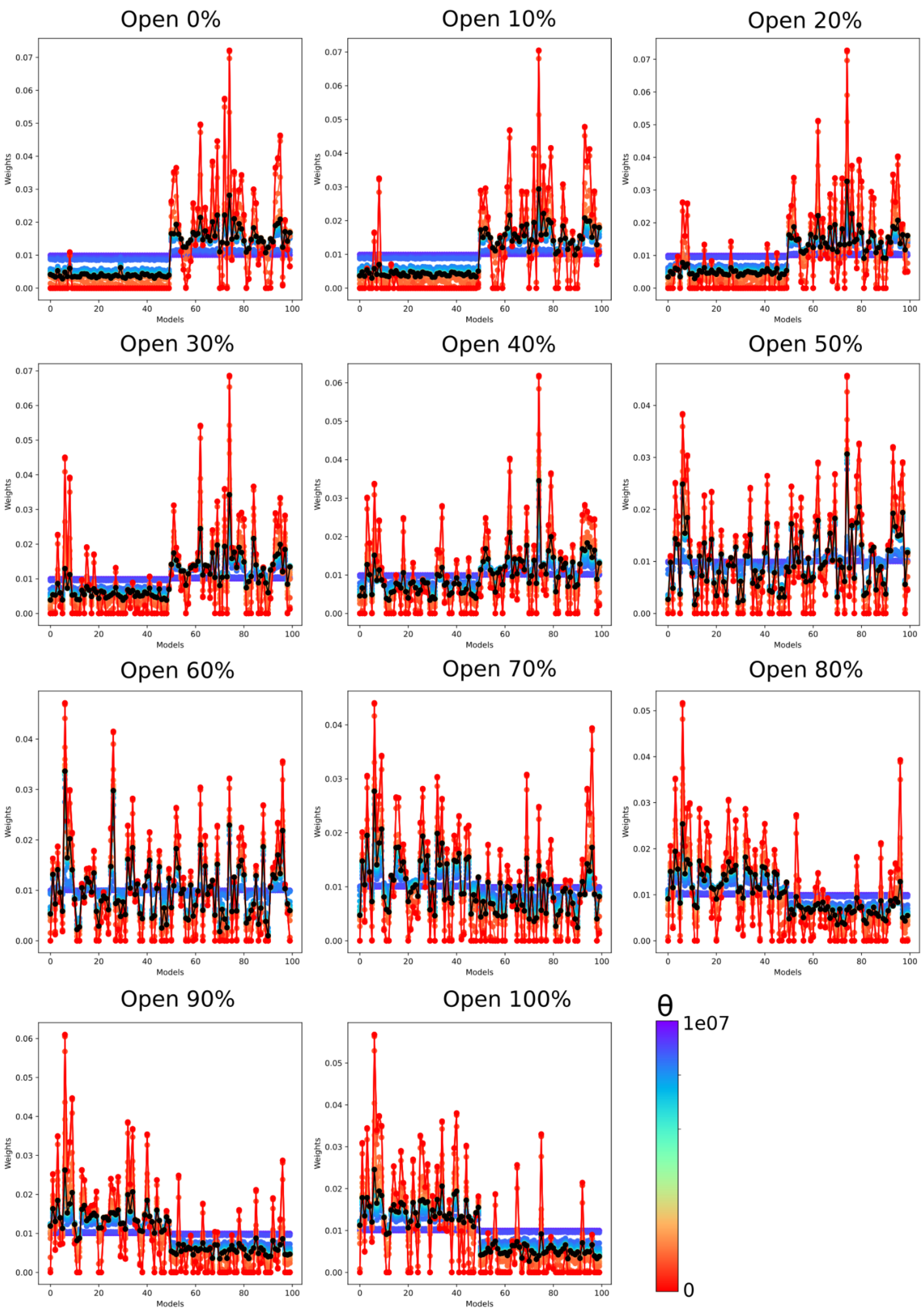


**Supplementary Figure 11**

The weights assigned to each model from the structural ensemble after the reweighting with the use of a 3 Å reference map with the different populations of the open state and a 1% noise level. The first 50 models represent the open conformation, while the last 50 models represent the closed conformation. Weights are presented for various θ values, with the optimal θ value shown in black.

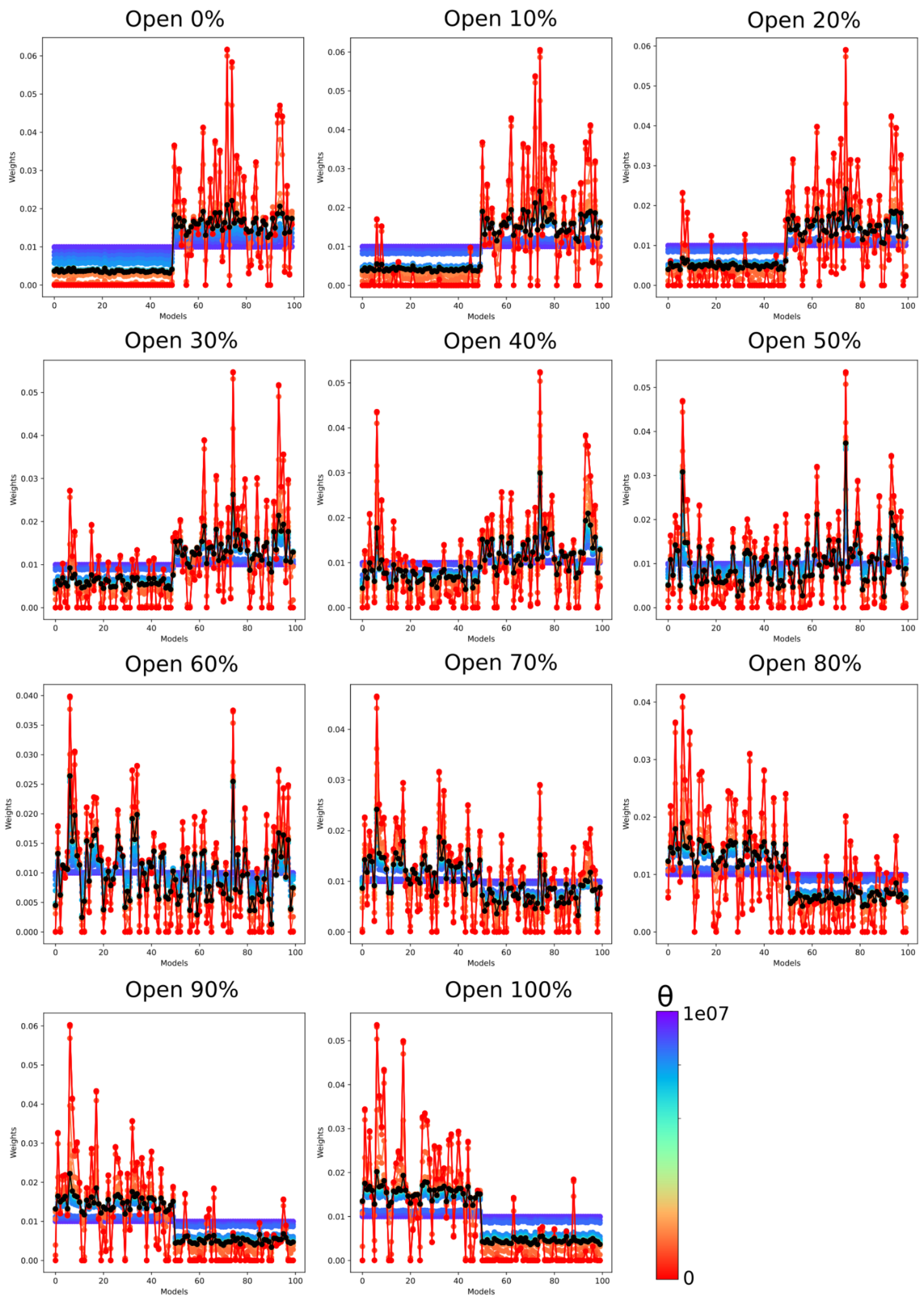


**Supplementary Figure 12**

The weights assigned to each model from the structural ensemble after the reweighting with the use of a 6 Å reference map with the different populations of the open state and a 10% noise level. The first 50 models represent the open conformation, while the last 50 models represent the closed conformation. Weights are presented for various θ values, with the optimal θ value shown in black.

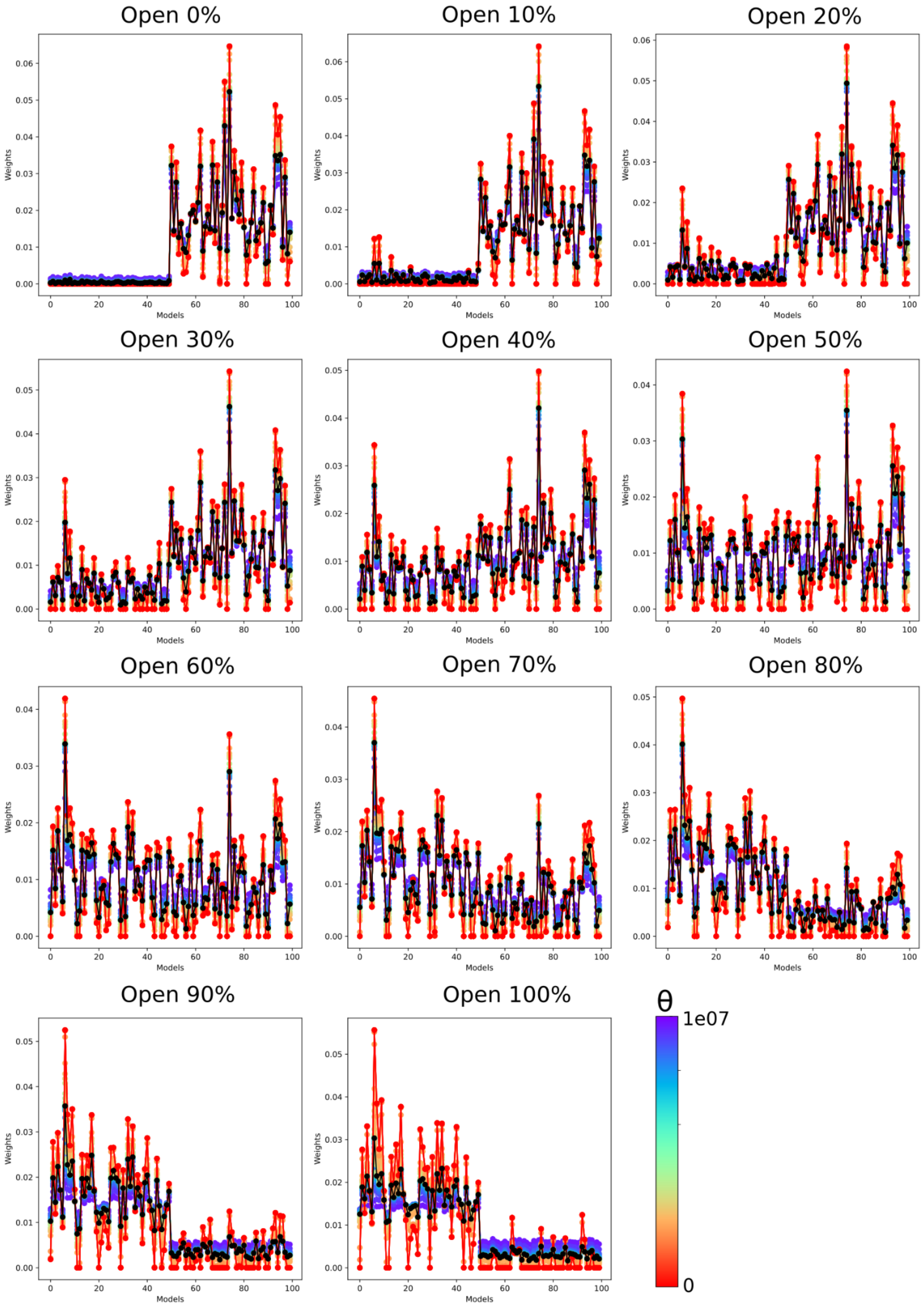


**Supplementary Figure 13**

The weights assigned to each model from the structural ensemble after the reweighting with the use of a 6 Å reference map with the different populations of the open state and a 1% noise level. The first 50 models represent the open conformation, while the last 50 models represent the closed conformation. Weights are presented for various θ values, with the optimal θ value shown in black.

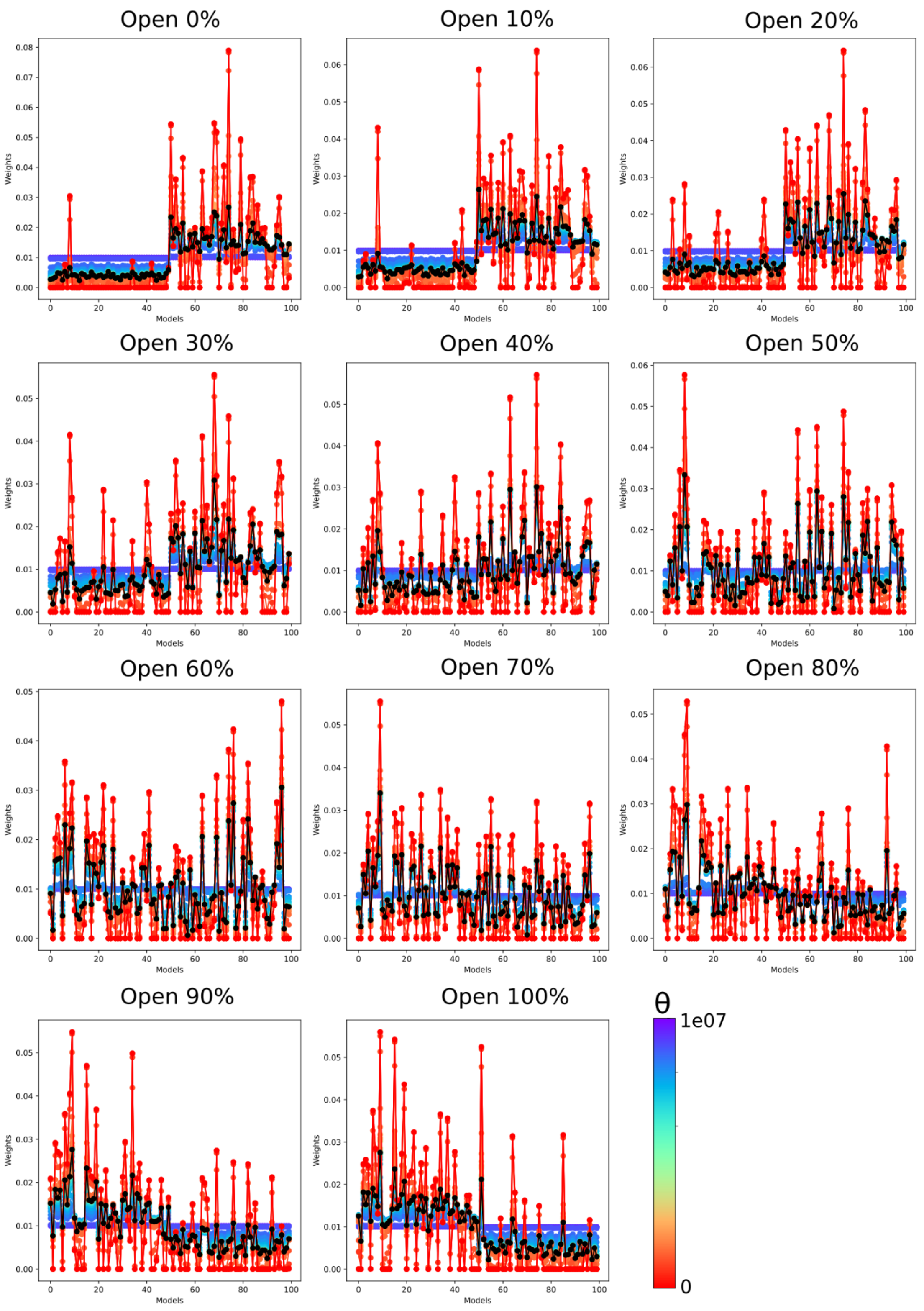


**Supplementary Figure 14**

The weights assigned to each model from the structural ensemble after the reweighting with the use of a 10 Å reference map with the different populations of the open state and a 10% noise level. The first 50 models represent the open conformation, while the last 50 models represent the closed conformation. Weights are presented for various θ values, with the optimal θ value shown in black.

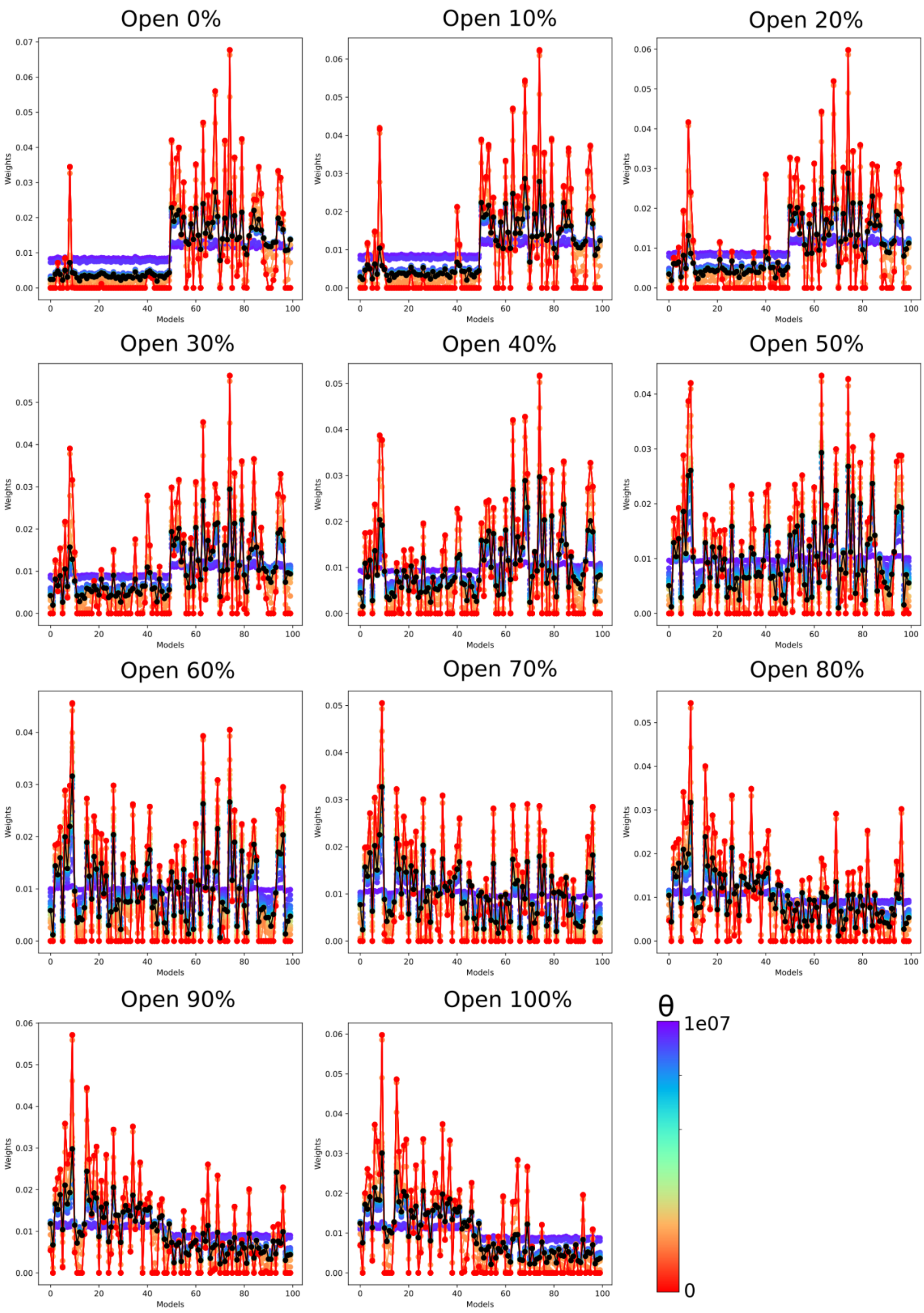


**Supplementary Figure 15**

The weights assigned to each model from the structural ensemble after the reweighting with the use of a 10 Å reference map with the different populations of the open state and a 1% noise level. The first 50 models represent the open conformation, while the last 50 models represent the closed conformation. Weights are presented for various θ values, with the optimal θ value shown in black.

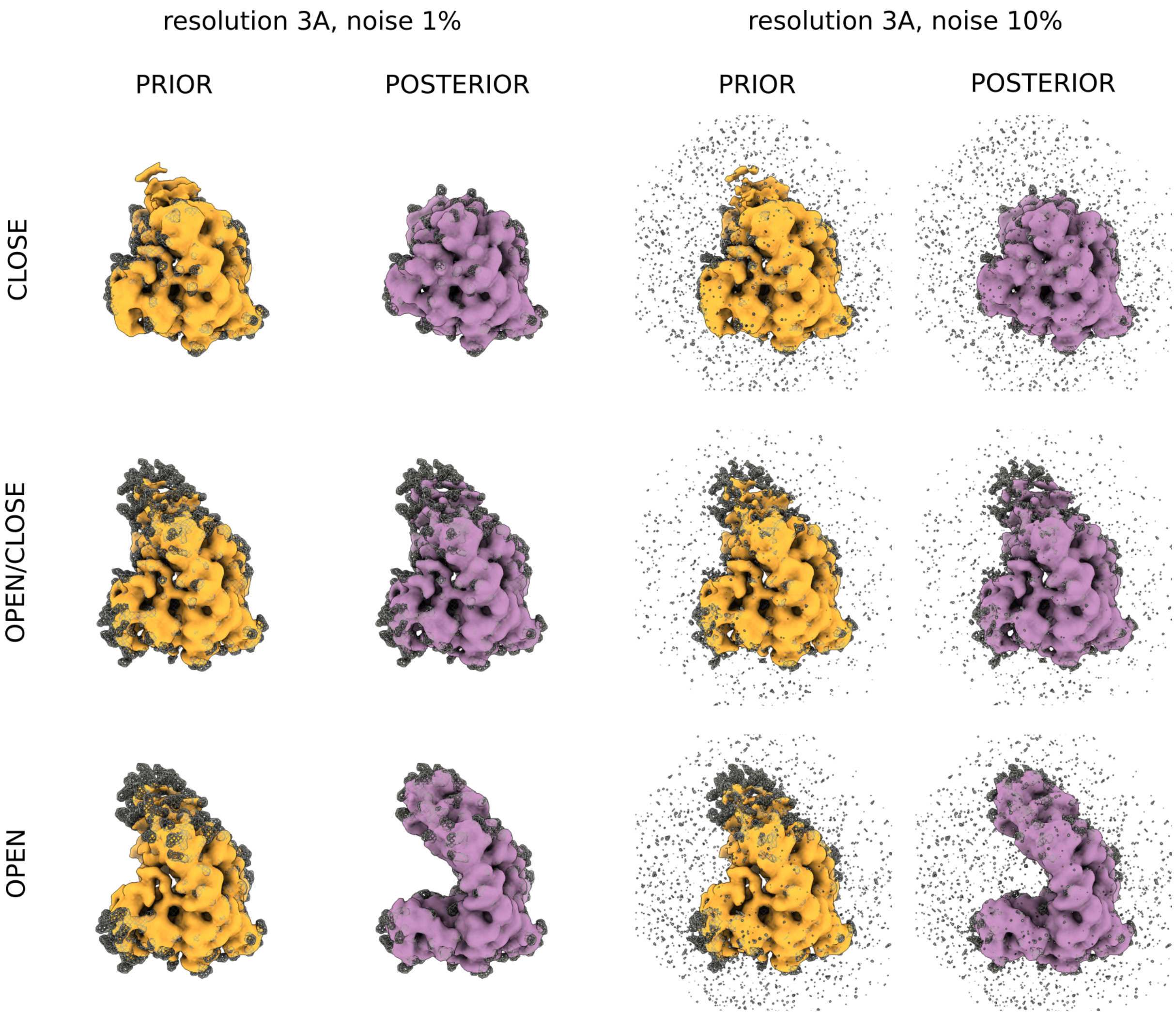


**Supplementary Figure 16**

Visual comparison of the 3 Å reference maps (displayed in grey mesh) with maps generated from the structural ensemble prior (in orange) and posterior (in purple) reweighting. The threshold for map visualisation is set at two standard deviations of the map density.

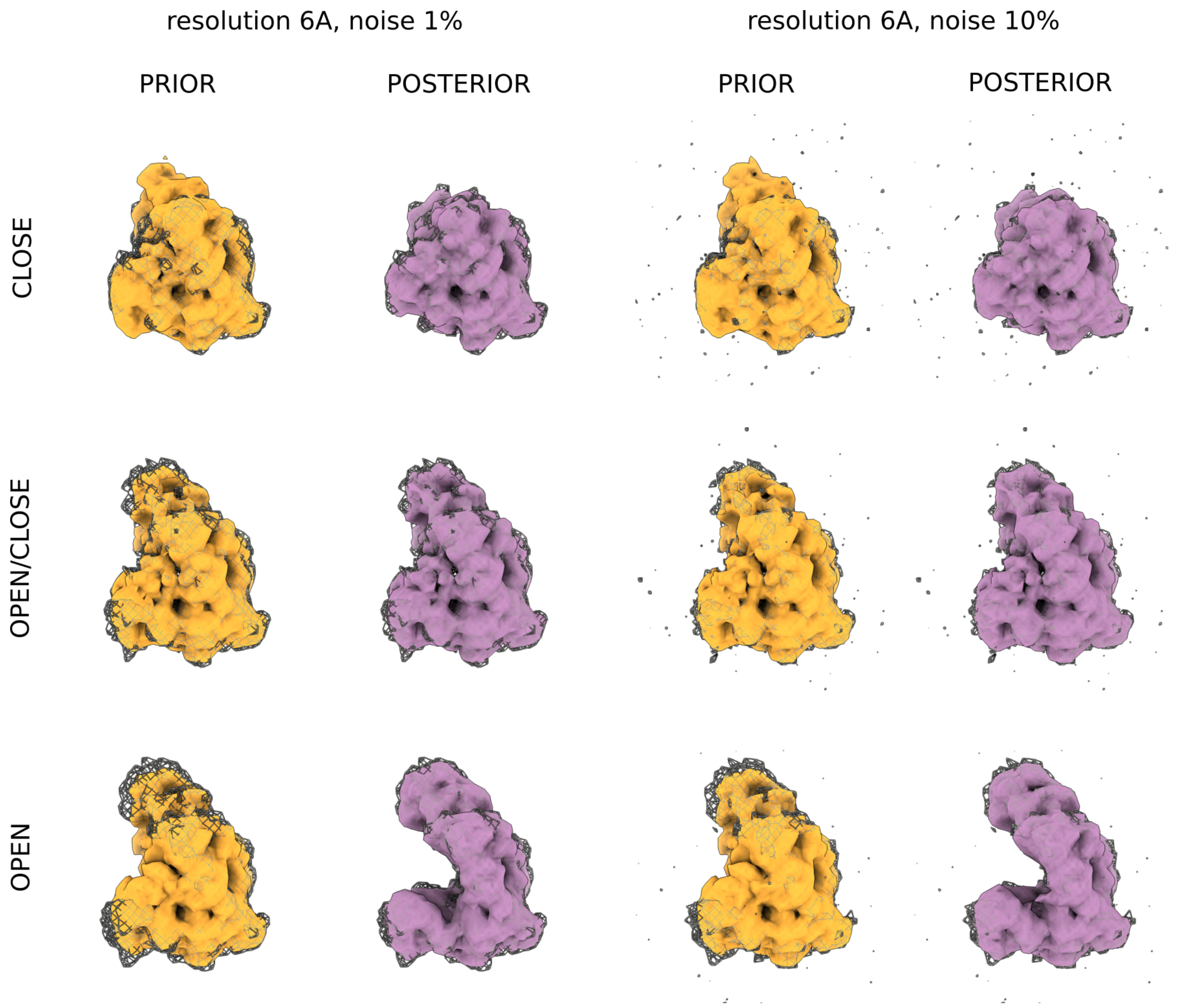


**Supplementary Figure 17**

Visual comparison of the 6 Å reference maps (displayed in grey mesh) with maps generated from the structural ensemble prior (in orange) and posterior (in purple) reweighting. The threshold for map visualisation is set at two standard deviations of the map density.

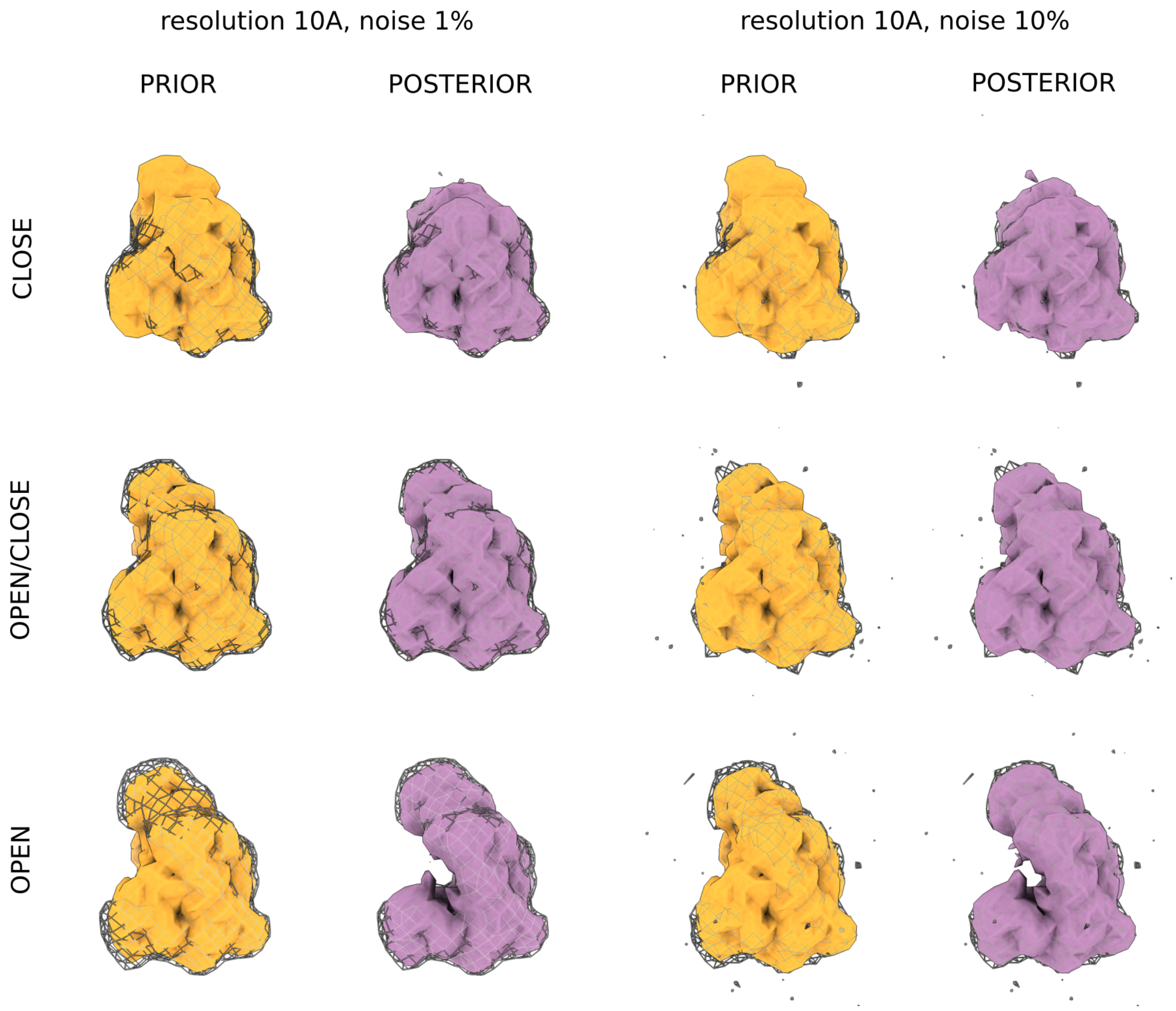


**Supplementary Figure 18**

Visual comparison of the 10 Å reference maps (displayed in grey mesh) with maps generated from the structural ensemble prior (in orange) and posterior (in purple) reweighting. The threshold for map visualisation is set at two standard deviations of the map density.

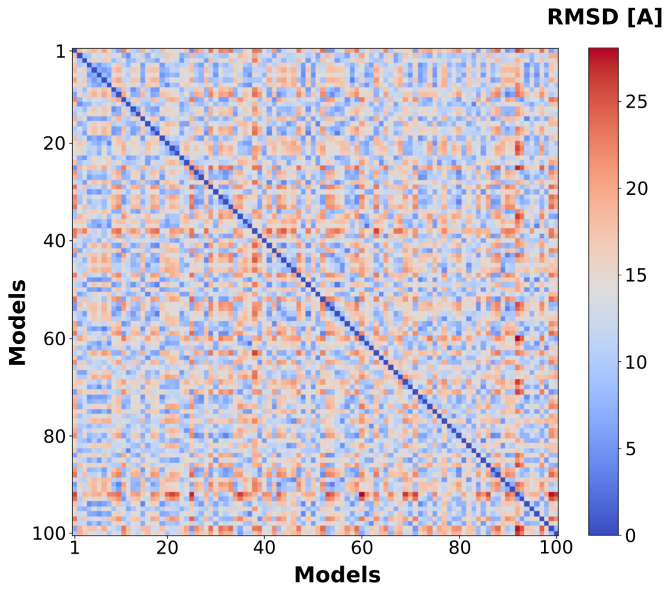


### Supplementary Figure 19

The RMSD matrix calculated for the FLN5 structural ensemble without structure superimposition.

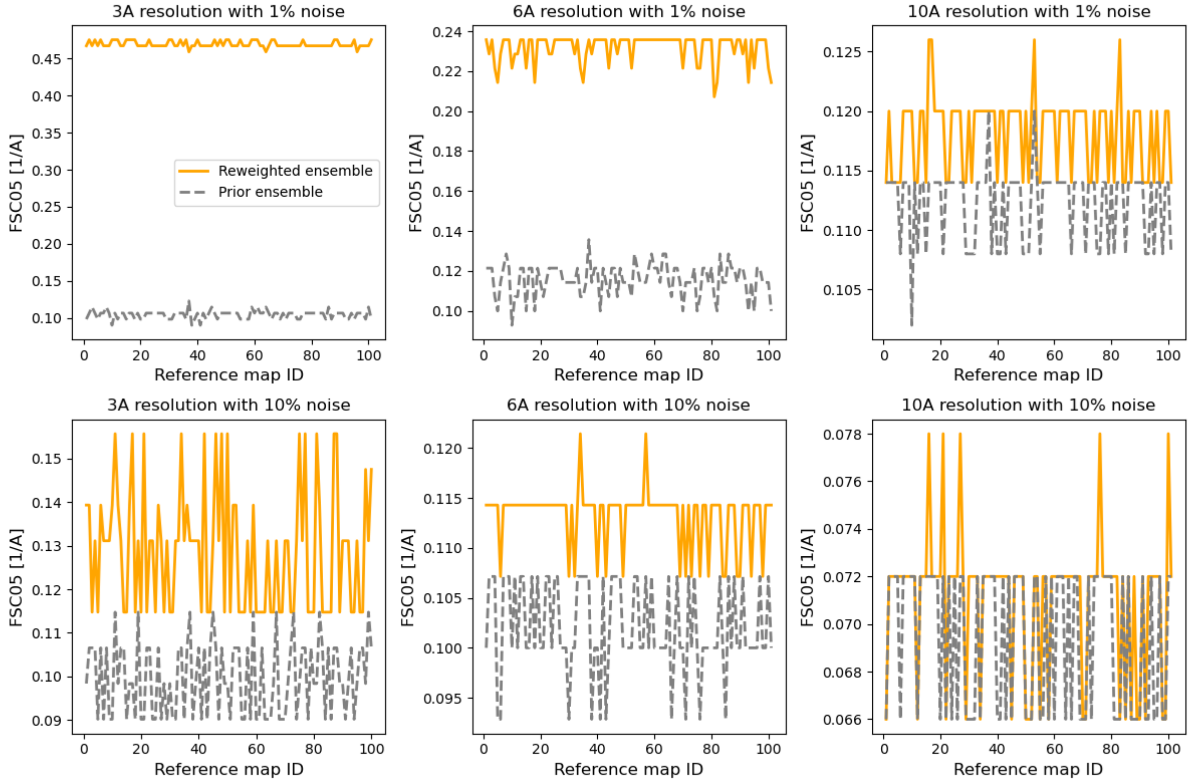


### Supplementary Figure 20

The FSC05 calculated between the reference map and the maps generated from the structural ensemble upon reweighting for each FLN5 dataset. The datasets varied in resolution and noise level.

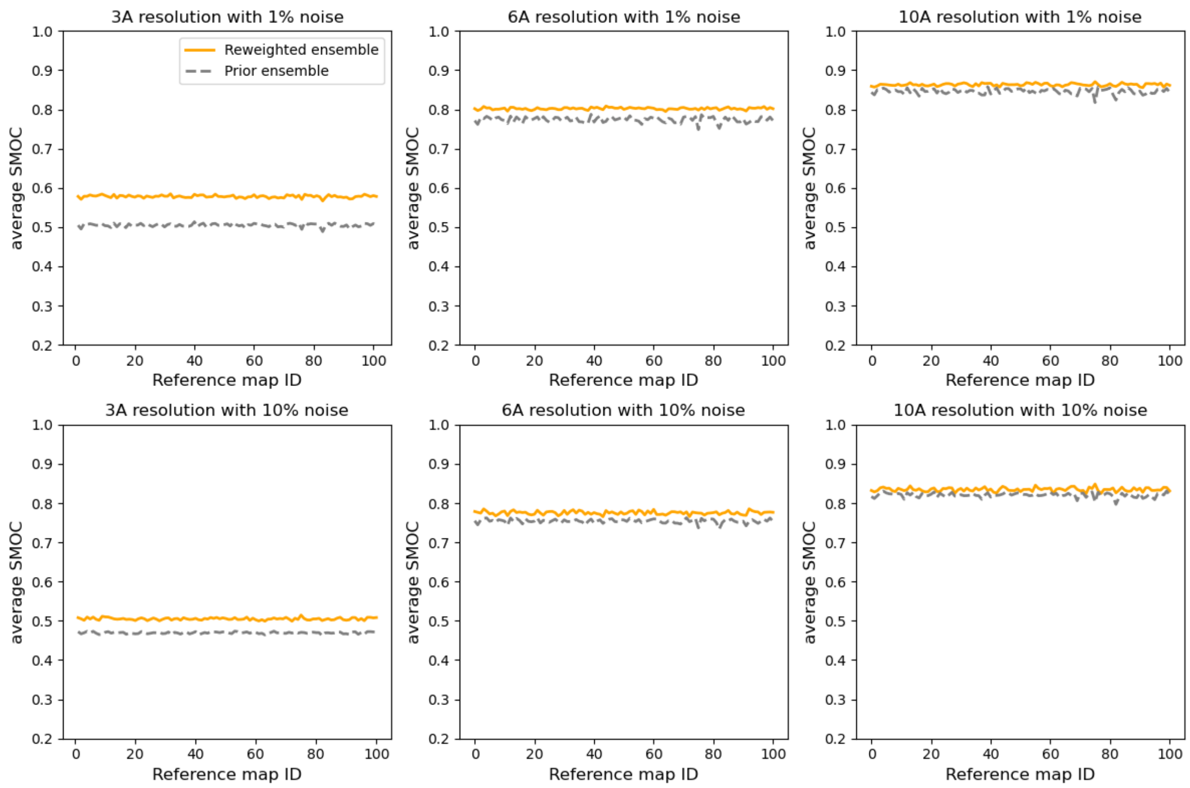


**Supplementary Figure 21**

The average SMOC score calculated between the reference map and the maps generated from the structural ensemble upon reweighting for each FLN5 dataset. The datasets varied in resolution and noise level.

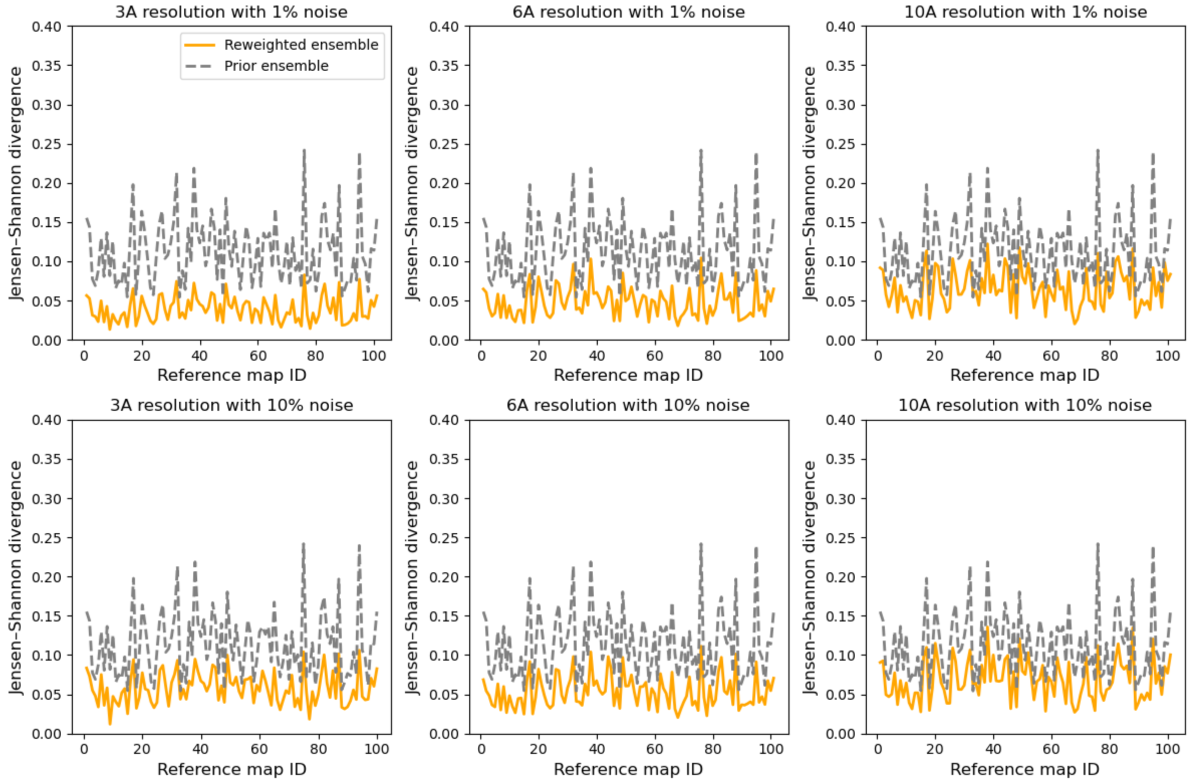


**Supplementary Figure 22**

The Jensen-Shannon divergence calculated between the MD ensemble (before and after reweighting) and the ensemble consisting of 10 structures used for map generation.

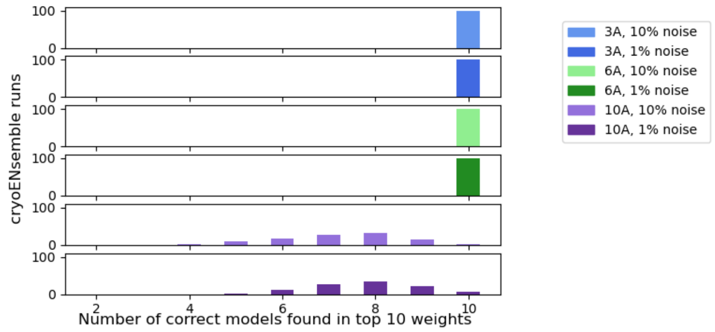


**Supplementary Figure 23**

Histograms presenting the number of correct models that were found in the top 10 models ranked by weights in cryoENsemble runs for each NC dataset.

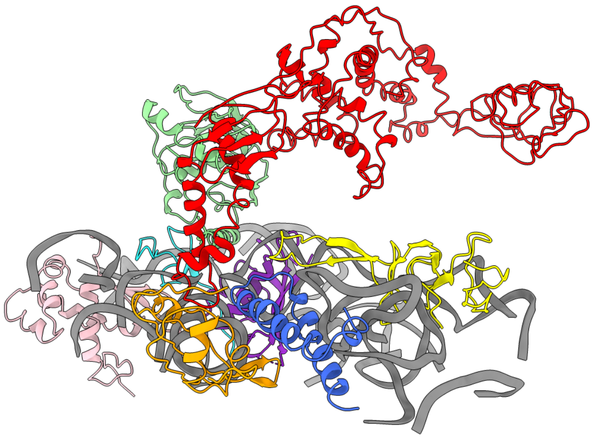

**Supplementary Figure 24**

The molecular system obtained from PDB id: 7D80 for all-atom structure-based model MD simulation that encompasses part of the 70S ribosome surface including rRNA (in grey), uL24 (in yellow), uL29 (in light blue), uL23 (in orange), uL17 (in pink), uL32 (in cyan), uL22 (in violet) ribosomal proteins as well as trigger factor (in red) and peptide deformylase (in light green).
